# Sequencing of methylase-accessible regions in integral circular extrachromosomal DNA reveals differences in chromatin structure

**DOI:** 10.1101/2021.03.31.437970

**Authors:** Weitian Chen, Zhe Weng, Zhe Xie, Yeming Xie, Chen Zhang, Zhichao Chen, Fengying Ruan, Juan Wang, Yuxin Sun, Yitong Fang, Mei Guo, Yiqin Tong, Yaning Li, Chong Tang

**Affiliations:** College of Life Sciences, University of Chinese Academy of Sciences, Beijing 100049, China; BGI Genomics, BGI-Shenzhen, Shenzhen 518083, China; Department of Biology, Cell Biology and Physiology, University of Copenhagen 13, 2100 Copenhagen,Denmark; Nantong University, Nantong, China, 226000; Nephrosis Precision Medicine Innovation Center, University of Beihua School of Medicine, Jilin 132011, China

**Author notes:** These authors contributed equally to this work (Weitian Chen, Zhe Weng). Correspondence: Chong Tang, Director of technology, BGI Shenzhen, China.

**Keywords:** ecDNAs, chromatin accessibility, methylation, m^6^A, methyltransferase

## Abstract

Although extrachromosomal DNA (ecDNA) has been intensively studied for several decades, the mechanisms underlying its tumorigenic effects have been revealed only recently. In the majority of conventional sequencing studies, the high-throughput short-read sequencing largely ignores the epigenetic status of most ecDNA regions except for the junctional areas. Here, we developed the sequencing of enzyme-accessible chromatin in circular DNA (CCDA-seq) method, which uses methylase to label open chromatin without fragmentation and exonuclease to enrich the ecDNA sequencing depth, followed by long-read nanopore sequencing. Using CCDA-seq, we observed significantly different patterns in nucleosome/regulator binding in ecDNA at a single-molecule resolution. These results deepen the understanding of ecDNA regulatory mechanisms.

## Introduction

Finding a cure for cancer has been a challenge for various reasons, such as oncogene amplification, tumor evolution, and genetic heterogeneity (Gillies et al. 2012; Ray Chaudhuri et al. 2016; Turajlic and Swanton 2017). These phenomena have been studied for many years. Recently, it has been demonstrated that circular extrachromosomal DNA (ecDNA) plays a critical role in carcinogenesis, as it promotes oncogene amplification (Wu et al. 2019), drives tumor evolution, and contributes to genetic heterogeneity (Turner et al. 2017; Paulsen et al. 2018b; Verhaak et al. 2019b). The circular ecDNAs is the DNA that is arranged next to chromatin in a circular structure, consisting of the featured head-to-tail junctional sequence and distal homologous genome sequence. The cancer-specific ecDNA may have an average size of 1.3 MB (Chiu et al. 2020). Although ecDNA was discovered since 1964 (Paulsen et al. 2018a), the elucidation of its role has been slow due to the lack of adequate molecular analytical techniques (Bailey et al. 2020).

The development of new techniques, including computational advances, enabled genetic and epigenetic studies of ecDNA, and attempts have been made to identify ecDNA from sequencing data using improved algorithms from specificity and sensitivity. Most algorithms, such as Circle-Map (Prada-Luengo et al. 2019), AmpliconArchitect (Deshpande et al. 2019), and CIRC_finder (Kumar et al. 2020), relied on the detection of the ecDNA junction sequence and enabled ecDNA identification in numerous cancer tissues (Turner et al. 2017; Paulsen et al. 2019; Verhaak et al. 2019a; Koche et al. 2020; Kumar et al. 2020), aging cells (Hull et al. 2019), plasma (Kumar et al. 2017; Zhu et al. 2017), and healthy somatic tissues (Møller et al. 2018b). However, due to the rareness of ecDNA in sequencing data, these approaches require the enrichment of ecDNA molecules; for example, circular DNA is obtained by the digestion of linear DNA with nucleases, followed by rolling circle amplification (Møller 2020). To further improve the accuracy of ecDNA detection, the long-read sequencing technology has been used to verify the ecDNA junction structure (deCarvalho et al. 2018) (Mehta et al. 2020). However, functional epigenetic studies of ecDNAs are currently lacking. Given the increasing awareness of ecDNA and its role in oncogene expression, understanding ecDNA chromatin state and transcription status is essential. The most recent and advanced theory proposed by Wu et al. offers insights into highly accessible chromatin region and high expression of oncogenes located within these regions in ecDNA using the assay for transposase-accessible chromatin sequencing (ATAC-seq), and Chromatin Immunoprecipitation Sequencing (Chip-seq) (Wu et al. 2019). Moreover, few studies have examined ecDNA epigenome, because it is difficult to analyze ecDNA junction structure and epigenome information simultaneously.

Given that ecDNA possesses a unique junction sequence pattern, its epigenetic information can be revealed by locating the neighboring junction regions without considering the distal region, which is thought to be indistinguishable from the linear genome sequences. However, there is a need for distal coverage of the ecDNA epigenome owing to the limitations due to the short-read sequencing and short fragmentation required for ATAC-seq (Buenrostro et al. 2015) and MNase-seq (Schones et al. 2008). Our research was enlightened by the existing long-read sequencing methods for assessing chromatin state, such as nanoNOMe-seq (Lee et al. 2020), SMAC-seq (Shipony et al. 2020), and fiber-seq (Stergachis et al. 2020). We used the N6-methyladenosine (m^6^A) methyltransferase EcoGII to soft label accessible chromatin regions without fragmentation and named this method sequencing of enzyme-accessible chromatin in circular DN<u>A</u> (CCDA-seq). Using this method, we enriched ecDNA by digesting the linear genome using nuclease. Nanopore sequencing accurately detected the m^6^A-probed ecDNA regions of accessible chromatin and junctional structure properties simultaneously in the long range. Using CCDA-seq, we found a high diversity of ecDNA regions of accessible chromatin and their coordination with distal regulators at a single-molecule resolution, which has not been reported before.

## Results

### CCDA-seq comprehensively maps accessible chromatin and nucleosome positioning in ecDNA at a multikilobase scale

ecDNA plays an important role in tumorigenesis due to the high accessibility of its chromatin and carried oncogenes (Wu et al. 2019). Conventional approaches to study chromatin accessibility are based on the concept that the chromatin protects the bound sequence from attack by transposase (Figure 1A) or MNase (Schones et al. 2008). In ATAC-seq, the open, accessible genome region is first preferentially tagged using transposase, followed by next-generation sequencing (NGS) (Figure 1A). However, this method is not employed in most integral ecDNA chromatin studies due to the homologous ecDNA/genome sequences, making the distinction between ecDNA and linear genome DNA difficult. In general, previous studies on ecDNA chromatin based on NGS of short reads only observed the chromatin status in the junction region (200 bp around the junction) and did not fully consider other distal ecDNA areas because of limitations of the techniques used (>200 bp to junction regions) (Figure 1A). To solve these problems, we built a generalized framework based on the concept of the SMAC-seq (Shipony et al. 2020) and fiber-seq (Stergachis et al. 2020). We applied soft labeling with the m^6^A methyltransferase EcoGII that preferentially methylates the adenosine in the openly accessible DNA region without fragmentation by a transposase (Figure 1A). To improve the ecDNA capturing efficiency, the exonuclease was introduced to remove the linear genome DNA (Gaubatz and Flores 1990). The integral ecDNA was sequenced by nanopore sequencing and the probed m^6^A was detected (Shipony et al. 2020). By analysis of the generated data, we first identified ecDNA molecules by head-to-tail junction locations and by dynamically mapping the segments of sequences to the genome (Figure 1B). Based on the head-to-tail junction locations, we then reassembled the partial ecDNA sequences as the new reference and identified the m^6^A signal based on the reassembled ecDNA sequence to prevent signal bias in the junction region (Figure 1B).

**Figure 1.**
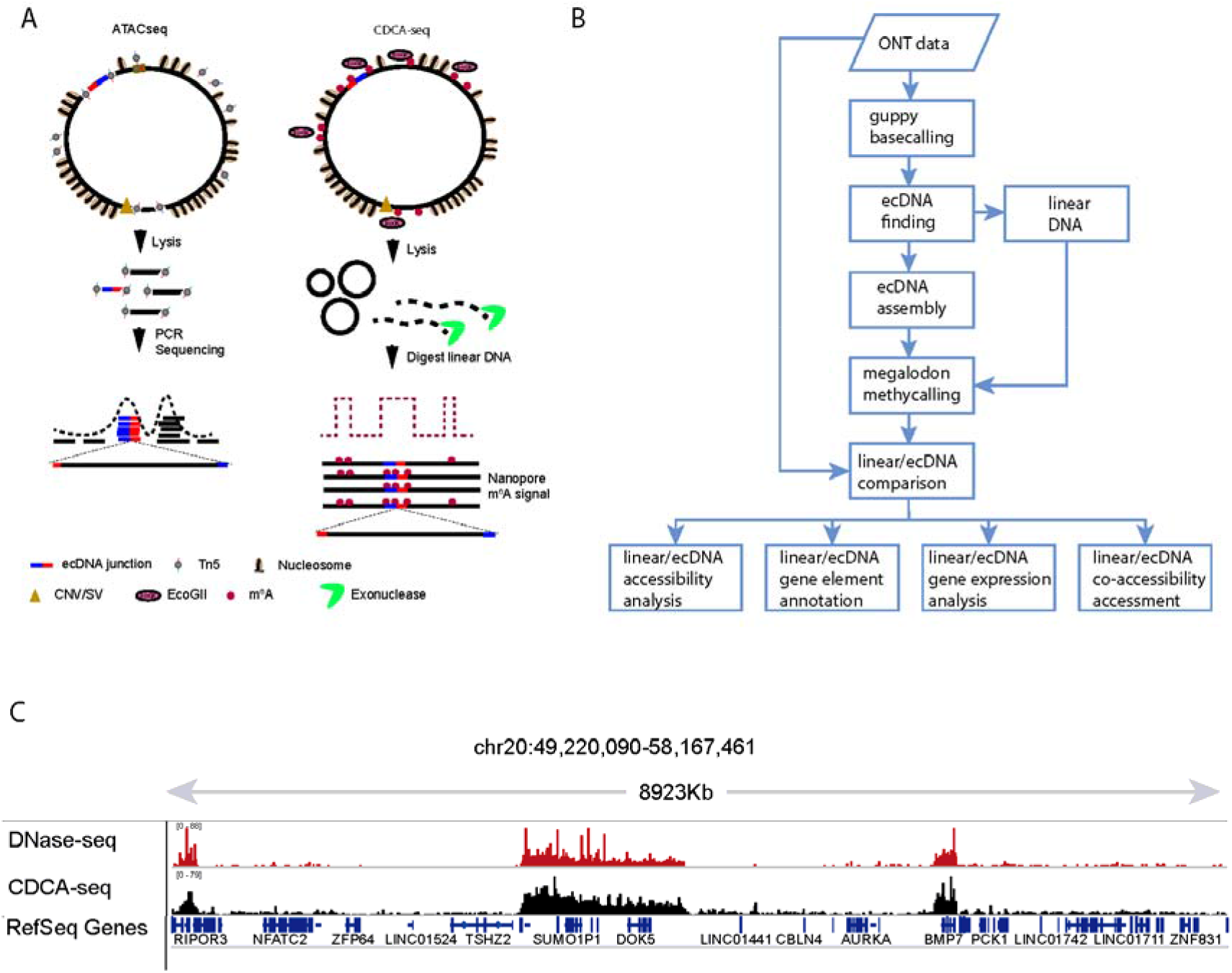
The CCDA-seq for profiling chromatin accessibility and nucleosome position at ecDNAs. (A) Intact chromatin is treated with m^6^A methyltransferase (EcoGII), which preferentially methylate DNA bases in open chromatin region on ecDNAs and linear DNAs. High molecular weight DNA is then isolated and subjected to exonuclease digestion to remove partially linear DNAs. The rest DNAs are subjected to nanopore library construction and nanopore sequencing. The data were aligned to genome to identify ecDNAs based on head-to-tail pattern. The methylated bases are used to reconstruct nucleosomes in ecDNAs and rest linear DNAs. In contrast, the ATAC-seq used the transposon to attack the open chromatin. The tagmentated short fragments are amplified and subjected to NGS. The short reads are aligned with genome to identify ecDNAs based. The mapped reads are calling as peaks to represent the open chromatin region. (B) The bioinformatics pipeline of CCDA-seq. The signal data were processed through guppy basecalling to generate sequence. The sequences were aligned to genome to identify the linear DNA and ecDNAs. We assembled the ecDNA sequence reference. Based on the ecDNA and linear DNA reference, we used Megalodon to call the m^6^A sites base on ecDNA and linear DNAs. Then we perform the accessibility analysis, gene element annotation, gene expression analysis, co-accessibility assessment. (C) Large aggregate CCDA-seq signal enrichments match closely with DNase-seq accessibility peaks. (Chr20:49,220,090-58,167,461)

The statistical analysis showed that the read length was between 10 and 100 kb, which is 50× broader than the junctional region observed in conventional ATAC-seq (Wu et al. 2019) (Supplemental Figure 1). The long-read feature also makes the nanopore sequencing method optimal for applications such as structure variation (SV), copy number variation (CNV), and ecDNA identification with better sensitivity and specificity (Huddleston et al. 2017). As expected, 80% of ecDNA molecules detected in our CCDA-seq could be validated through PCR (Supplemental Figure 2). ecDNA and residual linear DNA accounted, respectively, for 0.9% and 99.1% of the total sequencing reads (Supplemental Figure 3) after exonuclease treatment. The m^6^A probability distribution in Megalodon showed two distinct peaks for the treated sample. The distribution of the narrow peak with lower m^6^A probability (mean = 0.49) was similar to the background noise distribution (Supplemental Figure 4). Therefore, we set m^6^A methylation probability over 0.53 as the cutoff for the true m^6^A signal (Supplemental Figure 4). The real positive cutoff value was set as 0.53, and the m^6^A calling specificity and sensitivity were 0.99 and 0.92, respectively (Supplemental Figure 4). The residual linear DNA was used as internal control for validation using published ATAC-seq data (He et al. 2012). CCDA-seq achieved consistency and coherence with ATAC-seq data in various resolutions (Figure 1C, Supplemental Figure 5). Good concordance was also found when comparing our results with those obtained by other published methods (Shipony et al. 2020; Stergachis et al. 2020). The m^6^A labeling deviation was reverse proportional to the m^6^A ratio and was strongly reduced to 0.015 in m^6^A enriched region (Supplemental Figure 6). The impact of the exonuclease treatment and reproducibility have been also investigated (Supplemental Figure 7). These characteristics of CCDA-seq are critical for effectively measuring the accessibility of chromatin in linear and circular DNA molecules in the multikilobase range.

**Figure 2.**
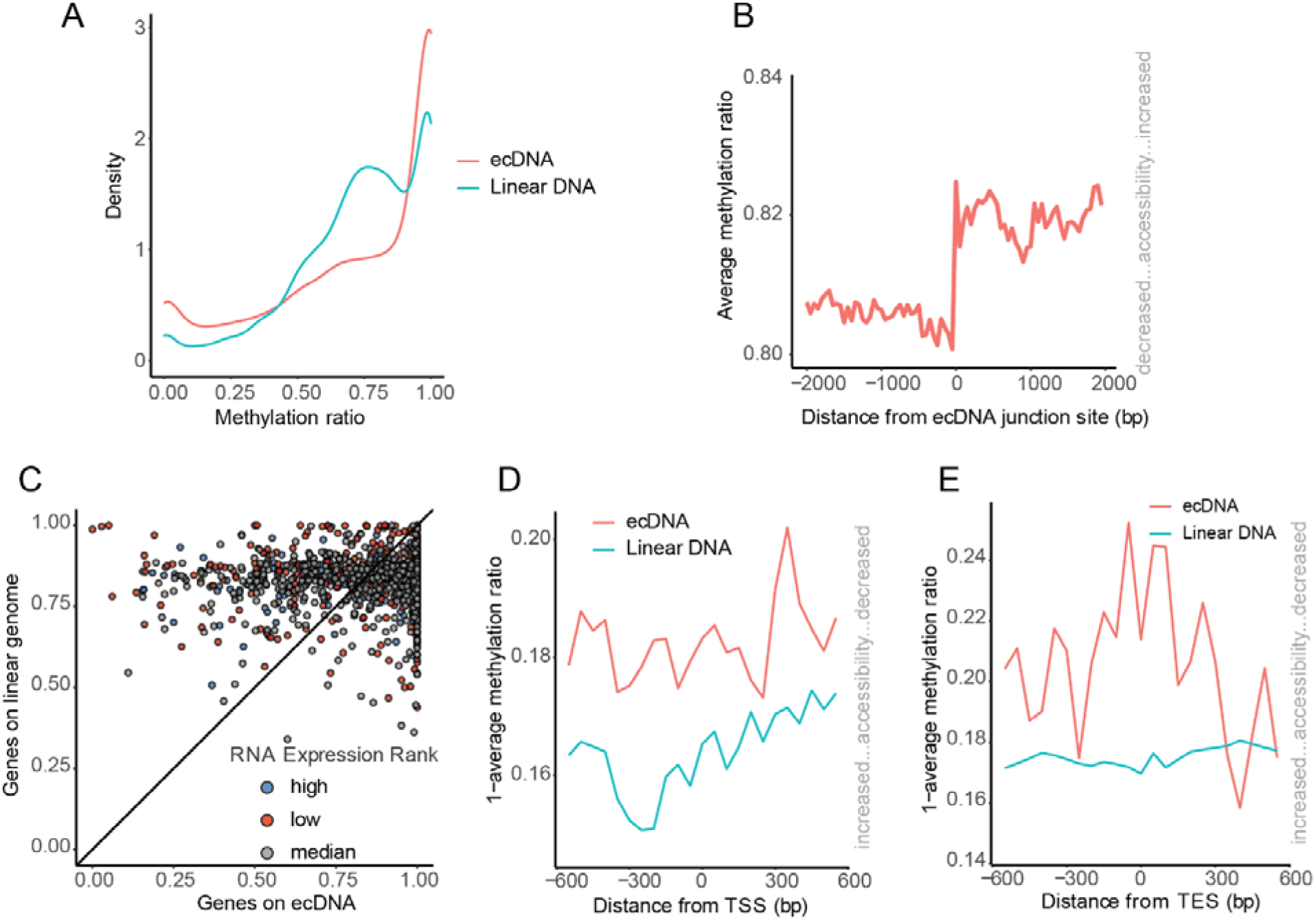
The ecDNA and linear DNA have the different chromatin accessibility pattern. (A) The density distribution of the methylation ratio on ecDNAs and linear DNAs. (B) The average chromatin status around ecDNA neighboring regions. The junction site and its right neighboring regions demonstrate the more open chromatin. (C) The average methylations of gene regions on ecDNA and linear DNAs (from TSS to TES). The genes were classified as two groups: I. The genes on linear DNA have more open chromatin structure than ecDNA carried genes (above centerline); II. The ecDNA carried genes have more open chromatin structure than the genes on linear DNA (below centerline). (D) Average CCDA-seq profile around transcription start site on ecDNAs and linear DNAs. (E) Average CCDA-seq profile around transcription end site on ecDNAs and linear DNAs. (aggregated over 50-bp windows sliding every 5 bp; the sequencing depth is normalized for ecDNA and linear DNA;see Methods for details)

**Figure 3.**
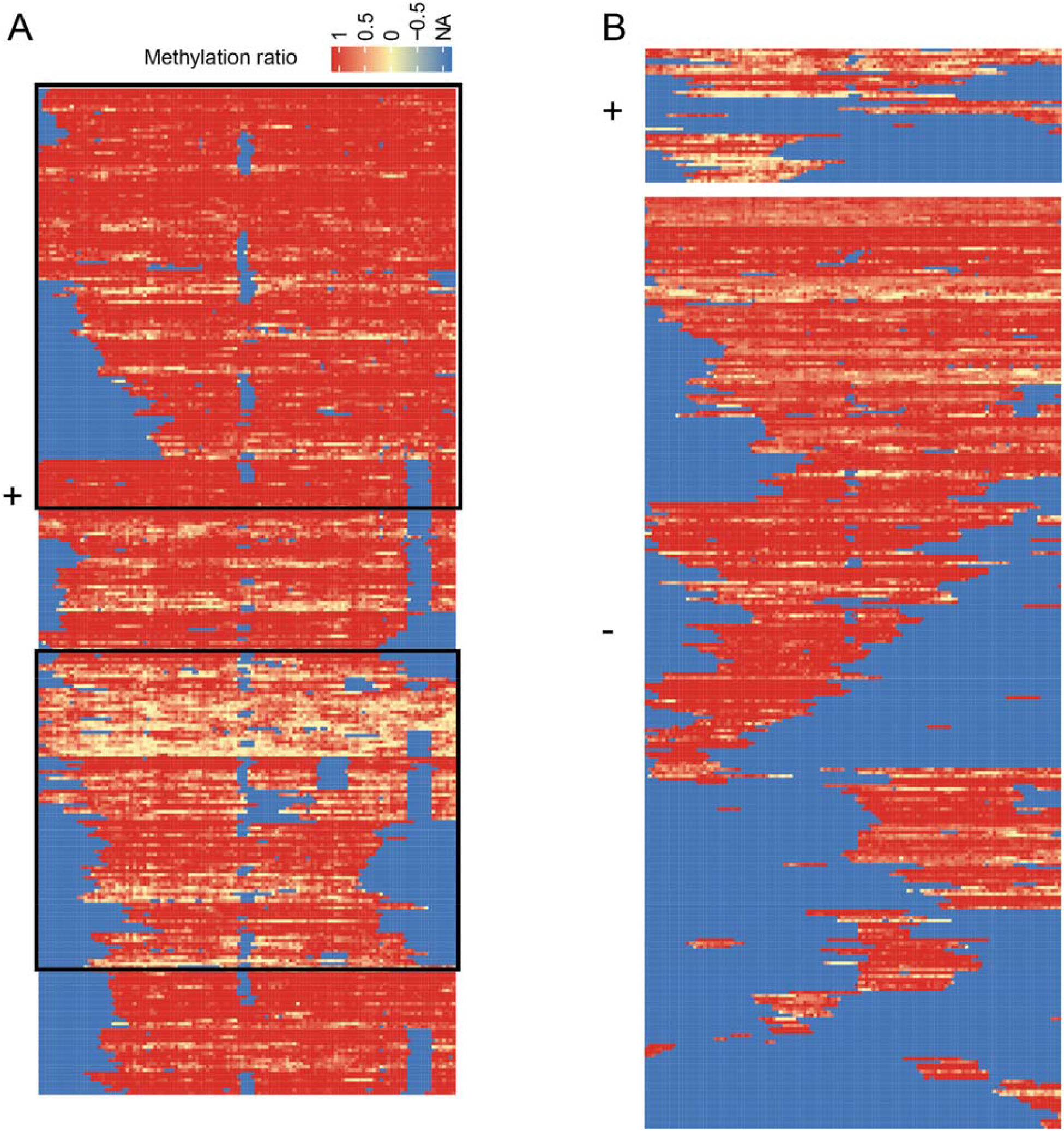
CCDA-seq reveals the distribution of alternative chromatin states of ecDNA arrays. A. Shown are all reads covering the linear DNA region chr10:42383201~42389201.The box highlights the active and inactive chromatins. B. Shown are all reads covering the ecDNA region chr10:42383201~42389201. The upper panel indicates the positive strand, and the lower panel indicates the negative strand.

**Figure 4.**
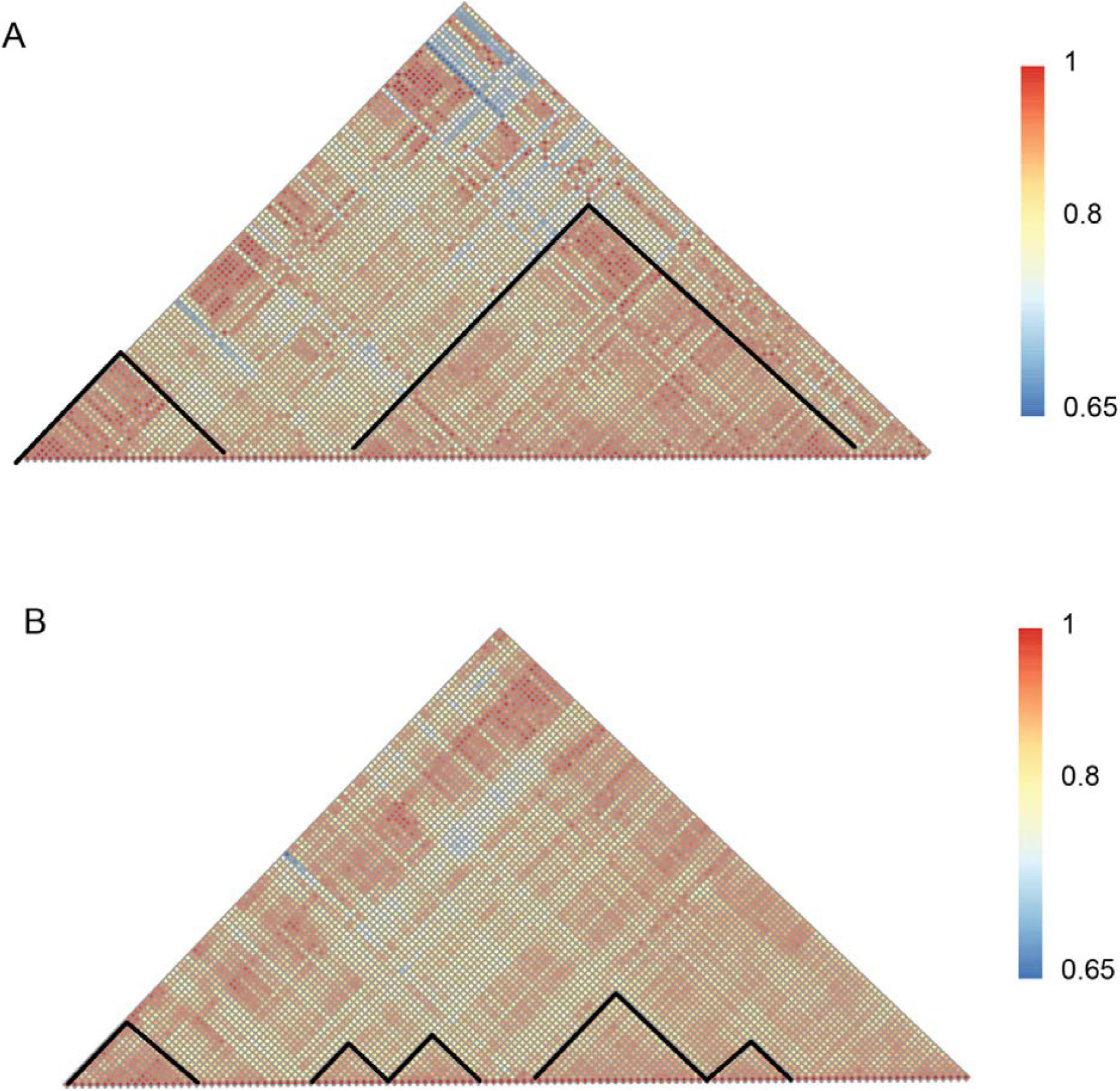
Chromatin co-accessibility profiles for the chr10:42383201~42389201 show correlation and low correlation in ecDNA and linear DNA. (A) Chromatin co-accessibility profiles for the chr10:42383201~42389201 show correlation and low correlation on ecDNA. Red indicates the positive correlation and blue indicates the low correlation. (B) Chromatin co-accessibility profiles for the chr10:42383201~42389201 show correlation and low correlation on linear DNA.

Another remarkable feature of CCDA-seq is that it enables single-molecule resolution of the ecDNA chromatin status. At the single-molecule level, the single base m^6^A probability varied from 0.6 to 1. (Supplemental Figure 8). In practice, the resolution of accessible chromatin regions was around 200 bp. We adopted a Bayesian procedure to aggregate methylation probabilities and derived the accurate single-molecule accessibility calls over windows of arbitrary size (Supplemental Figure 8). In summary, CCDA-seq offers attractive features in terms of elucidation of the integral ecDNA chromatin status in the multikilobase range at a single-molecule resolution.

### Diverse patterns of ecDNA chromatin accessibility

Evidence from other studies obtained by ATAC-seq and Chip-seq suggests that the active chromatin status and highly accessible ecDNA chromatin may be associated with high levels of oncogene transcription (Wu et al. 2019). To distinguish the ecDNAs molecules from the linear DNA molecules in ATAC-seq and Chip-seq, it is necessary to screen out the short reads (~200 bp) spanning the non-homologous end-joining ecDNA sequence. One problem with these approaches is the potential bias to neglect the distal regions due to focusing on the ~200 bp reads neighboring ecDNA junctional sequences. CCDA-seq, as a long-read technology, may facilitate precise ecDNA detection (Huddleston et al. 2017) (Møller et al. 2018a) (deCarvalho et al. 2018) and observation of the distal chromatin status in integral ecDNA. We obtained an extensive catalog of 12,997 different ecDNA molecules formed from chromosomal breakpoints between 0.05 kb and up to 100 kb (Supplemental Table 1). Gene ontology (GO) analysis of the genes harbored by these ecDNA molecules revealed significant enrichment in the GO terms GTPase-related activity, channel activity, and nucleoside-triphosphatase activity, i.e., processes playing essential roles in cancer progression (Supplemental Figure 9) (House et al. 2015; Kazanietz and Caloca 2017). RNA-seq data analysis showed that there were 340 highly expressed ecDNA genes (25% rank), 464 moderately expressed genes (25~75% rank), and 589 genes with low expression (75%–100% rank), indicating that not all ecDNA genes are highly expressed.

By comparing the average chromatin accessibility between ecDNA and homologous linear DNA, we found that the ecDNA chromatin is twofold more accessible than that the linear DNA chromatin (Figure 2A). These findings reinforce the general notion that ecDNA amplification results in higher oncogene transcription (Wu et al. 2019), coupled with the enhanced chromatin accessibility in the junctional region. The CCDA-seq data were subjected to the detailed mapping of the ecDNA chromatin status. We found that chromatin in the ecDNA junctional areas is significantly more accessible than in other linear homologous regions (Figure 2B). This is an interesting finding, as it suggests that the conclusions drawn by observing only the junctional areas after the conventional ATAC-seq may be biased and not necessarily relate to the whole ecDNA chromatin. We calculated the average fractions of m^6^A methylation from the gene transcription start site (TSS) to the gene transcription end site (TES) on each gene-spanning read. A pairwise scatter plot of the average accessibility between ecDNA genes and linear genome genes showed that 63% of gene regions are more accessible in the ecDNA than in the linear DNA (Figure 2C). Comparing the ecDNA and linear DNA chromatin profiles around the TSS/TES (+/− 500 bp) revealed a significant difference in nucleosome depletion/occupancy patterns (Figure 2D, E). The nucleosome organization may impact access to ecDNA (Figure 2D, E). Considering that 63% of gene regions were more accessible on the ecDNA than on the linear DNA, we further plotted the chromatin structure around TSS/TES (+/− 500 bp) of these genes (Supplemental Figure 10). The formation of nucleosome depletion regions (NDRs) on linear DNA is restricted to 200 bp before TSSs. In contrast, the NDRs on ecDNA are distributed uniformly (Supplemental Figure 10). The other 37% of gene regions are more accessible on the linear DNA than on the ecDNA. The TSSs/TESs (+/− 500 bp) were also significantly more accessible on the linear DNA than on the ecDNA with different NDR patterns (Supplemental Figure 11). The formation of large NDRs was restricted to TSSs on the linear DNA, which was not observed on the ecDNA.

Another illustration of the complex interplay between chromatin states in the ecDNA and linear DNA relates to the transcriptional activity. The chromatin of the linear DNA active genes (top 25% rank) is largely devoid of nucleosomes on TSSs due to the extremely high transcription activity (Supplemental Figure 12). In contrast, the chromatin structure of the ecDNA active genes adopts a distinct conformation, implying that ecDNA is regulated by different mechanisms (Supplemental Figure 12). For the transcriptionally inactive genes, the stationary nucleosome states are shown in the linear DNAs (Supplemental Figure 13). In contrast, ecDNA molecules still have the active nucleosome organization in the regions of 300 bp before TSSs, suggesting that chromatin accessibility is necessary but not sufficient for the enhancer or promoter activity in the ecDNA (Supplemental Figure 13). In conclusion, ecDNA and linear DNA have significantly different nucleosome depletion/occupancy patterns in various conditions, suggesting their distinct gene regulatory mechanisms.

### Chromatin status in the ecDNA and linear genome DNA at a single-molecule resolution

The conventional ATAC-seq is based on statically calling the peak of the enriched read in a specific region (Buenrostro et al. 2015). Recent single-molecule and single-cell accessibility measurements suggested that ATAC-seq of cell populations represent an ensemble average of distinct molecular states (Klemm et al. 2019). An essential attribute of the CCDA-seq is a possibility to determine ecDNA chromatin accessibility at a single-molecule resolution by taking the advantage of small variance (Supplemental Figure 6) and increased cumulative probability in segments (Supplemental Figure 8). Measuring chromatin accessibility of the single linear DNA has also been done in the SMAC-seq (Shipony et al. 2020) and fiber-seq (Stergachis et al. 2020).

We then asked whether CCDA-seq could reveal multiple chromatin accessibility states in ecDNA. The chromatin structure of the linear DNA (chr10: 42383201–42389251) adopts two distinct conformations: an inactive nucleosomal state and a state largely devoid of nucleosomes due to extremely high transcription activity (Conconi et al. 1989) (Figure 3A). It is thought that the majority of cancer cells exhibit active nucleosome status on ecDNAs. As expected, 70% of ecDNA molecules come from the very active chromatin state (Figure 3A). We observed highly heterogeneous nucleosome depletion/occupancy patterns in ecDNA, and most chromatin molecules were not very active in the positive strand, suggesting distinct transcriptional regulation of ecDNA (Figure 3B, upper panel). Some regulator enzymes may occupied the positive strand and restricted the chromatin accessibility. The highly active ecDNA chromatin were also observed in other regions (Supplemental Figure 14). To avoid a conclusion biased by the methylase heterogeneous activity, the other upstream and downstream regions were chosen as quality controls.

To further explore CCDA-seq resolution limits, we studied methylation patterns in more detail. We next quantified strand-specific DNA accessibility and observed a strand-asymmetric DNA accessibility pattern in the linear genome (Supplemental Figure 14). The strand-asymmetric DNA accessibility pattern was also observed in ecDNA, and both strands displayed high heterogeneity (Figure 3B, Supplemental Figure 14). This strand-specific heterogeneity in methylation potential within the nucleosome may inform about how transcription factors interact with nucleosome-associated DNA *in vivo*.

Wu et al. showed that ecDNA enables ultra-long-range chromatin contact, permitting distant interactions with regulatory elements (Wu et al. 2019). We next examined co-accessibility patterns in the ecDNA and linear genome DNA by assessing nucleosome positioning correlations. The nucleosomes have higher correlation values on the ecDNA than on the linear DNA (Figure 4A, B; Supplemental Figure 15). Moreover, ecDNA and linear DNA adopt significantly different chromatin co-accessibility patterns (Figure 4A, B; Supplemental Figure 15). Average co-accessibility profiles on the linear DNA revealed a detectable correlation between nucleosome positions up to two to three nucleosomes away. For the ecDNA, this correlation was further and up to 20 nucleosomes away (Figure 4A, B; Supplemental Figure 15). These results agree with the high-resolution chromosome conformation capture (HiC) result (Wu et al. 2019) in that the ecDNA is characterized by the distant chromatin interaction. It was interesting to note that ecDNA demonstrated some ultra-distant anticorrelated states. Overall, ecDNA molecules were highly heterogeneous and exhibited remote chromatin interactions, suggesting their different regulation mechanisms compared to those of linear DNAs.

## Discussion

Understanding of ecDNA functions may prove to be essential for the elucidation of tumorigenesis mechanisms (Turner et al. 2017; Paulsen et al. 2018b; Verhaak et al. 2019b). Many ecDNA molecules have been identified in various cancer tissues (Paulsen et al. 2019; Koche et al. 2020; Kumar et al. 2020) (Verhaak et al. 2019a) (Turner et al. 2017). There has been an increasing research focus on the status of ecDNA chromatin to resolve the problem of ecDNA oncogene amplification (Wu et al. 2019). However, most studies focused on short sequencing reads with junctional sequences detected to avoid the false-positive identification of ecDNA and to precisely determine the ecDNA epigenetic status. A large subgroup (60%) of the ecDNAs covered regions that are not unique in the reference genome, which complicated their identification (Moller et al. 2015). In this study, we used nanopore sequencing to evaluate integral ecDNA chromatin accessibility on ecDNA long strands by applying m^6^A methyltransferase to label open chromatin without fragmentation. Consistent with the previously reported findings (Wu et al. 2019), 63% of ecDNA molecules carried genes with more accessible chromatin structure than that of the linear DNA. However, in the remaining fraction of ecDNA (37%), chromatin of the gene regions was less accessible than in the corresponding linear DNA parts. Notably, the nucleosome depletion/occupancy patterns were significantly different between ecDNA and linear DNA. Our single-molecule resolution method allows footprinting of protein and nucleosome binding as well as determination of the epigenetic signature of chromatin accessibility. It is hoped that this study will contribute to more comprehensive understanding of the ecDNA epigenome regulation.

In our experiments, we treated DNA samples with an exonuclease that removed most of the linear DNA molecules and increased the sequencing depth for the ecDNA (0.9%). Some identified linear DNA molecules may be generated from the ecDNA homologous regions without junctions, but the likelihood of that was around 0.9%, which is negligible. Compared with the parameters in the non-digest direct sequencing, we only obtained 0.1% of ecDNA-related reads (Supplemental Figure 16). The circular eccDNA enrichment was 10×. The exonuclease treatment not only improved ecDNA sequencing coverage, but also ecDNA detection specificity (Supplemental Figure 2). However, DNA purification process could damage large-size ecDNA molecules over 1 Mb (Smith and Cantor 1989). Such damaged ecDNA could be digested during exonuclease digestion and missed in the sequencing. A method that gently purifies large DNA molecules would be preferable in further large-scale ecDNA studies.

Megalodon is the latest software (compared with Tombo), chosen for m^6^A signal calling. In the ecDNA m^6^A calling, Tombo ignored half of the sequences or lost most ecDNA molecules for unknown reasons (Supplemental Figure 17). The sensitivity of Tombo for ecDNA m6A signals was 83% less than that of Megalodon. Although Megalodon improved the sensitivity of ecDNA m^6^A calling, it did not address the issue of the false-positive m^6^A signal, that most adenosine bases could be recognized as m^6^A with a probability of 0.4–1 using Megalodon. The only known way to solve the false positive issue is to employ data training with negative control samples (Supplementary Figure 4). We used 0.53 as m^6^A probability cutoff, successfully discriminating the m^6^A and false-positive signals with sensitivity of 0.92 and specificity of 0.99. In general, Megalodon performed better in ecDNA analysis, and its specificity improved following data training.

In the sequencing data, we found that the methylated treated DNA generated more data than the non-methylated DNAs, which was not consistent with the SMAC-seq and fiber-seq data (Shipony et al. 2020) (Stergachis et al. 2020). The highly open chromatin with highly methylated sites may have been enriched using our method. In our laboratory experiments, we found that the heavily modified DNA was more resistant to exonuclease digestion, which led to the enrichment of modified DNA. The non-treated sample showed a lower overall methylation level (Supplemental Figure 18). However, the nucleosome occupancy positions were not significantly affected by the exonuclease treatment (Supplemental Figure 19). Moreover, in the strand-specific view, the reverse strand reads are generally less abundant than the positive strands. This may also be due to the different patterns of methylation of the positive and negative strands, which could result in different digestion efficiencies. This problem is usually overcome by increasing sequencing depth and using normalization methods. We also suggest sequencing both treated and non-treated samples for ecDNA sequencing coverage as this further improves quantification accuracy.

Only 63% of gene regions were within highly accessible chromatin in our experiments. However, Wu et al. showed using ATACsee technology that ecDNA molecules are mostly located in highly accessible chromatin (Wu et al. 2019). When comparing results from all other regions with the published data, a good agreement was found in that 80% of the areas were highly accessible in the ecDNA (Supplemental Figure 20). Most of the areas of highly accessible chromatin were distributed in the intron and intergenic regions (Supplemental Figure 21). The reasons for this remain unclear, but our results indicate that ecDNA has a highly open chromatin structure, especially in the intergenic and intronic regions.

CCDA-seq is useful for studying the chromatin status of integral ecDNA, offering deep insights into the distinct mechanisms of ecDNA regulation. However, the ecDNA enrichment step requires exonuclease treatment, causing the loss of mega ecDNAs. It is assumed that future advances will help address the problems of DNA damage during the purification and the insufficient sequencing depth. The CCDA-seq will help the scientific community to understand different mechanisms of ecDNA regulation, especially in cancer development.

## Method

### Cell culture

Human mammary gland carcinoma cell line MCF-7 was obtained from ATCC. MCF-7 were grown in DMEM(Gibco,11995065) supplemented with 10% FBS (Gibco,10099141), 0.01mg/ml insulin(), and 1% penicillin-streptomycin(Gibco, 15140122). The cell line was regularly checked for mycoplasma infection (Yeasen, 40612ES25).

### Nuclei isolation and MTase treatment

Cells were grown to 70-80% confluency, and were collected by TrypLE (Gibco,12604013). After 300xg centrifuge for 5 minutes, nuclei were isolated with lysis buffer (100 mM Tris–HCl, pH 7.4, 10 mM NaCl, 3 mM MgCl_2_, 0.1 mM EDTA, 0.5% CA630) for 5 minutes on ice. Nuclei were centrifuged at 300xg in wash buffer (100 mM Tris–HCl, pH 7.4, 10 mM NaCl, 3 mM MgCl_2_, 0.1 mM EDTA) at 4 degree, and washed twice for 5 minutes and counted. 1×10^6 intact nuclei were subjected to an m^6^A methylation reaction mixture containing 1x Cutsmart buffer (NEB), 200U of non-specific adenine methyltransferase M.EcoGII (NEB, M0603S), 300mM sucrose, and 96 uM S-adenosylmethionine (NEB, B9003S) in 500ul volume. The reaction mixture was set up at a 37-degree thermomixer with shaking at 1000rpm for 30 minutes. S-adenosylmethionine was replenished at 640uM every 7.5 minutes at 7.5, 15, and 22.5 minutes into the reaction mixture. The reaction was stopped by adding an equal volume of stop buffer (20 mM Tris-HCl pH 7.4, 600 mM NaCl, 1% SDS, 10 mM EDTA). No methylation controls were treated in the same conditions without adding M.EcoGII in the reaction mixture. The samples were then treated with 20ul of Proteinase K (20mg/ml) at 55 degrees overnight, and the DNA was extracted with phenol: chloroform extraction and ethanol precipitation.

### ecDNA isolation, purification, and sequencing

ecDNA was isolated by Circle-Seq(Møller 2020) method, which digested linear DNA with modifications. Briefly, 10ug of M.EcoGII treated DNA was subjected to a reaction mixture containing 1x plasmid-safe reaction buffer, 20U plasmid-safe ATP-dependent DNase (Lucigen, E3101K), 1mM ATP, and nuclease-free water was supplemented to a final volume of 100ul. The reaction mixture was incubated at 37 degrees for 7 days. Every 24 hours, the reaction mixture was replenished by adding 20U plasmid-safe ATP-dependent DNase, 1mM ATP, and 0.4ul 10X plasmid-safe reaction buffer. Digested ecDNA was purified with 1.8X AMpure XP beads (Beckman Coulter).

Purified ecDNA was prepared for nanopore sequencing by ligation kit LSK-SQK108(ONT). The samples were 10kb by Covaris G tubes, end-repaired and dA-tailed using NEBnext Ultra II end-repair module (NEB), followed by clean-up using 1.8X AMpure XP beads. Sequencing adaptors and motor proteins were ligated to end-repaired DNA fragments using blunt/TA ligase master mix (NEB), followed by clean-up using 0.4x AMpure XP beads. 1ug adaptor-ligated samples per flow cell were loaded onto PRO-002 flowcells and run on PromethION sequencers for up to 72h. Data were collected by MinKNOW v.1.14.

### Base-calling and Linear DNA methylation calling

Reads from the ONT data were processed using Megalodon V2.2.9, which used Guppy base caller to base-calling, and Guppy model config res_dna_r941_min_modbases-all-context_v001.cfg released into the Rerio repository was used to identify DNA m^6^A methylation. Megalodon_extras was used to get per read modified_bases from the Megalodon basecalls and mappings results. To further explore the accurate threshold of methylation probability, a control sample with almost no m^6^A methylation was used as background noise, and the Gaussian mixture model was used to fit the methylation probability distribution generated by Megalodon.

### ecDNA calling

ONT Reads meet the following conditions were defined as ecDNA molecules performed by the inner mappy/minimap2 aligner (Li 2018). (1) One segment (>1kb) of an ONT read was mapped to the genome at one site, and another segment (>1kb) was mapped to the genome at another site. (2) Two segments were mapped to the same chromosome. (3) Two segments were mapped to the same strand of the genome. (4) Two segments in a pair showed outward orientation.

### Nanopore ecDNA methylation calling

Due to ecDNA special structure, the m^6^A calling cannot be successfully performed by aligning to the reference genome, especially for junctional regions. The custom python script was used to assemble ecDNA reference genome sequences according to the table generated from the previous step. Considering that the read length might be longer than the ecDNA reference, the ecDNA reference was subsequently preprocessed by adding 10M N to the ends to increase the mapping efficiency. The downstream step is performed in a similar way as linear DNA methylation calling.

### Annotation and methylation configuration

TES, TTS, CDS, and other gene elements were downloaded from UCSC Table Browser, And the gene elements were processed into 50bp bin for downstream analysis. Linear DNA and ecDNA were also processed to the size of 50bp bin and sliding for 5bp. The accessibility score over multi base-pair windows was calculated as methylation ratio = m^6^A bases in all covered reads under bin/ adenosine bases in all covered reads under the bin.

### RNA-seq data analysis

The RNA-seq data of MCF-7 was downloaded from the Gene Expression Omnibus (GEO) repository database with the accession number GSE71862. The gene expression was divided into three categories: high, medium, and low, representing 25%, 25%-75%, and 75% gene expression rank, respectively.

### Co-accessibility assessment

To evaluate co-accessibility patterns along the genome, we applied COA as follows. Each chromosome in the genome was split into windows of size w. For each such window (i, i + w), we identified another window (j,j+w) such that the span (i,j,w) was covered by ≥N reads. For each single spanning molecule k, accessibility scores (A) in each bin were then aggregated and binarized as described above. The local co-accessibility matrix between two windows was calculated:

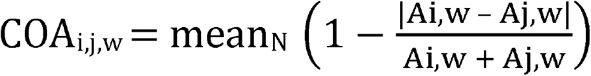

## Data availability

Nanopore raw data are available at China National GeneBank (CNGB) with project number of CNP0001299.

## Acknowledgment

Funding: This research was supported by the Science, Technology, and Innovation Commission of Shenzhen Municipality (grant number JSGG20170824152728492). The supporter had no role in designing the study, data collection, analysis, and interpretation, or in writing the manuscript.

## Author contributions

C.T. designed and supervised the experiments. Z.W. and X.Z perform the lab experiments; W.T.C. performs the bioinformatics data analysis. All others joined the data analysis.

## Competing interest

The authors declare no competing interests.

## Notes

### Competing Interest Statement

The authors have declared no competing interest.

