## Supplemental Figures for "Sequencing of methylase-accessible regions in integral circular extrachromosomal DNA reveals differences in chromatin structure"

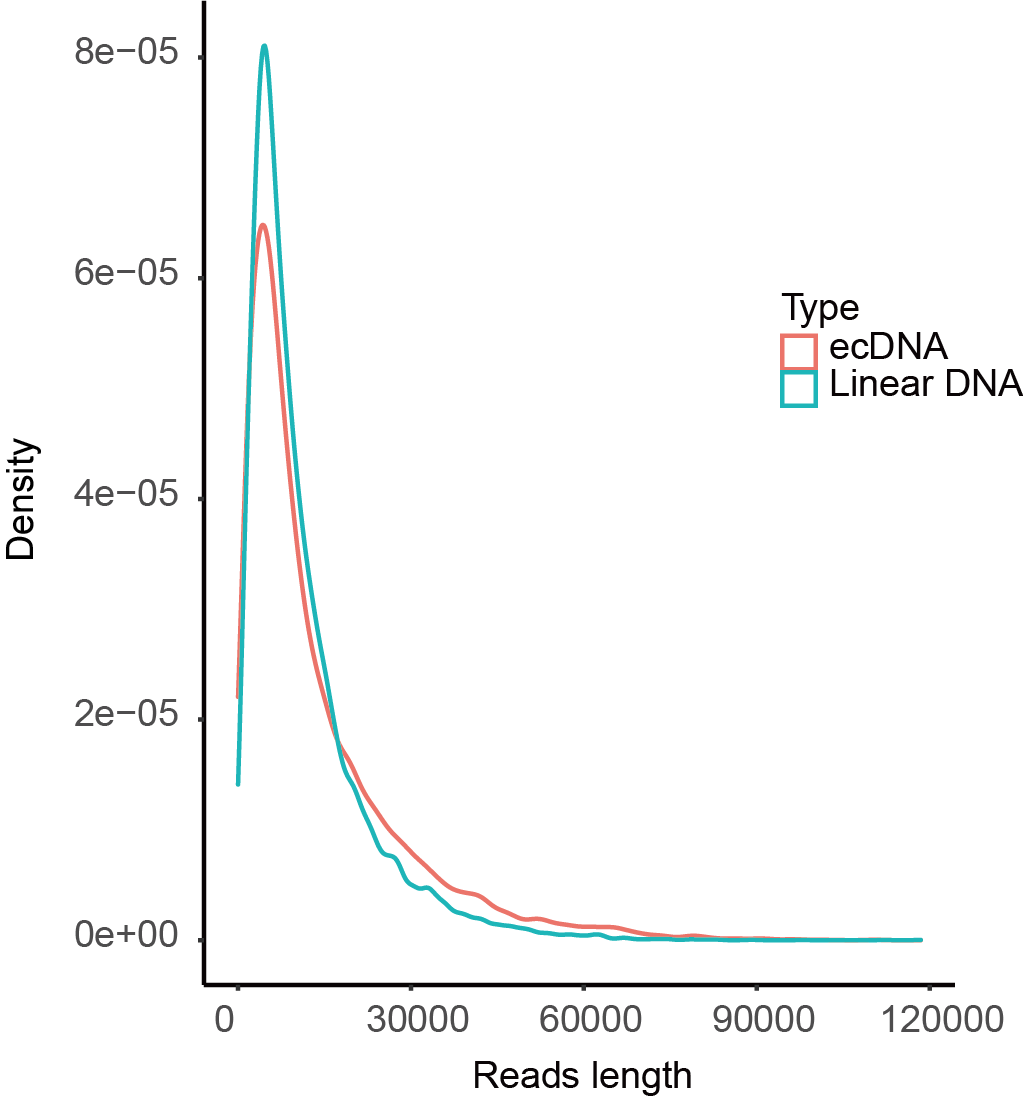

**Supplemental Figure 1. The length distribution of reads identified as ecDNAs or linear DNAs.**

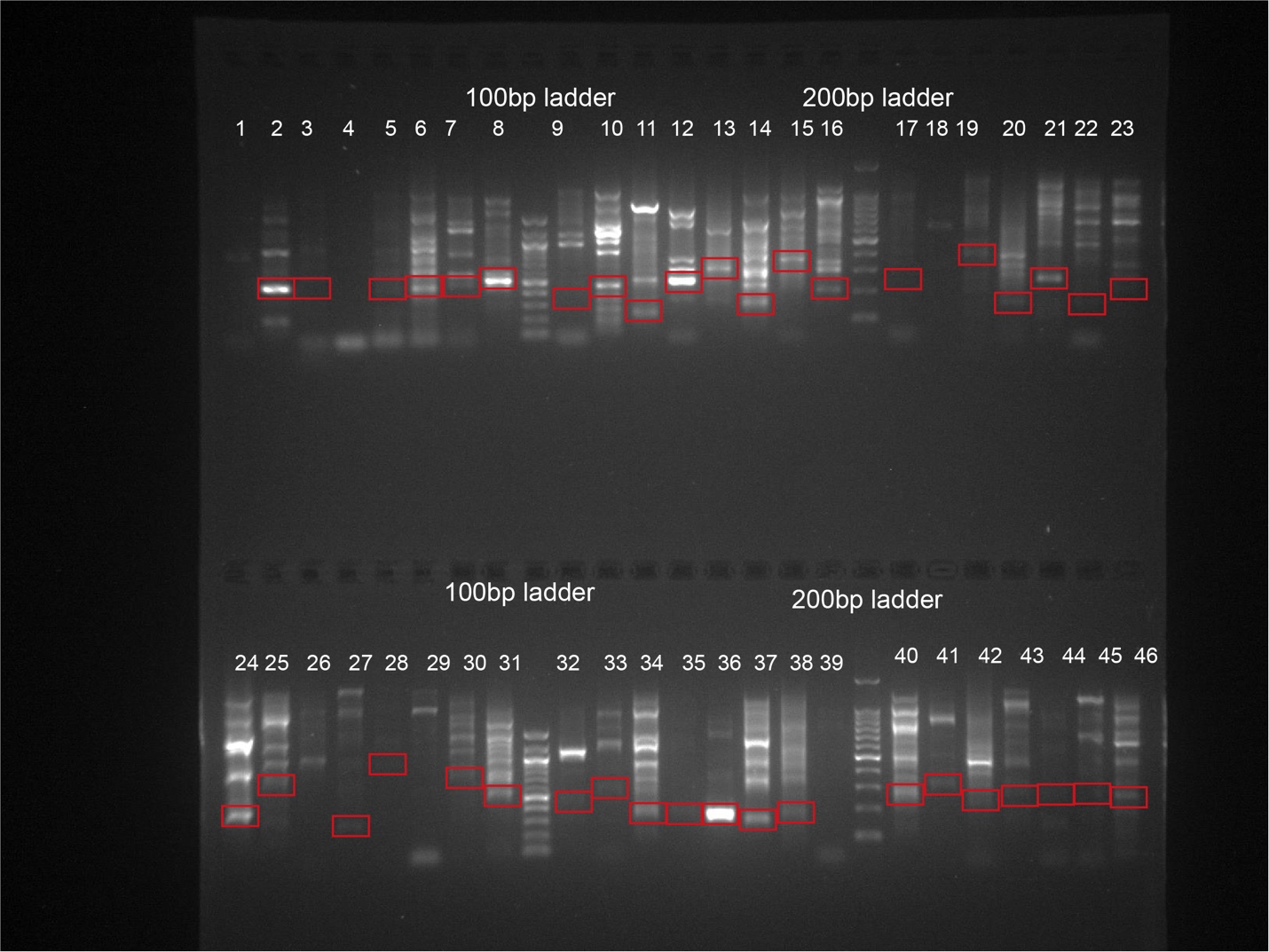

**Supplemental Figure 2. The PCR validation of the identified ecDNAs.** The red box indicates the expected DNA size. We found many other structure variations around the junction region, so the PCR targeted fragments are not unique.

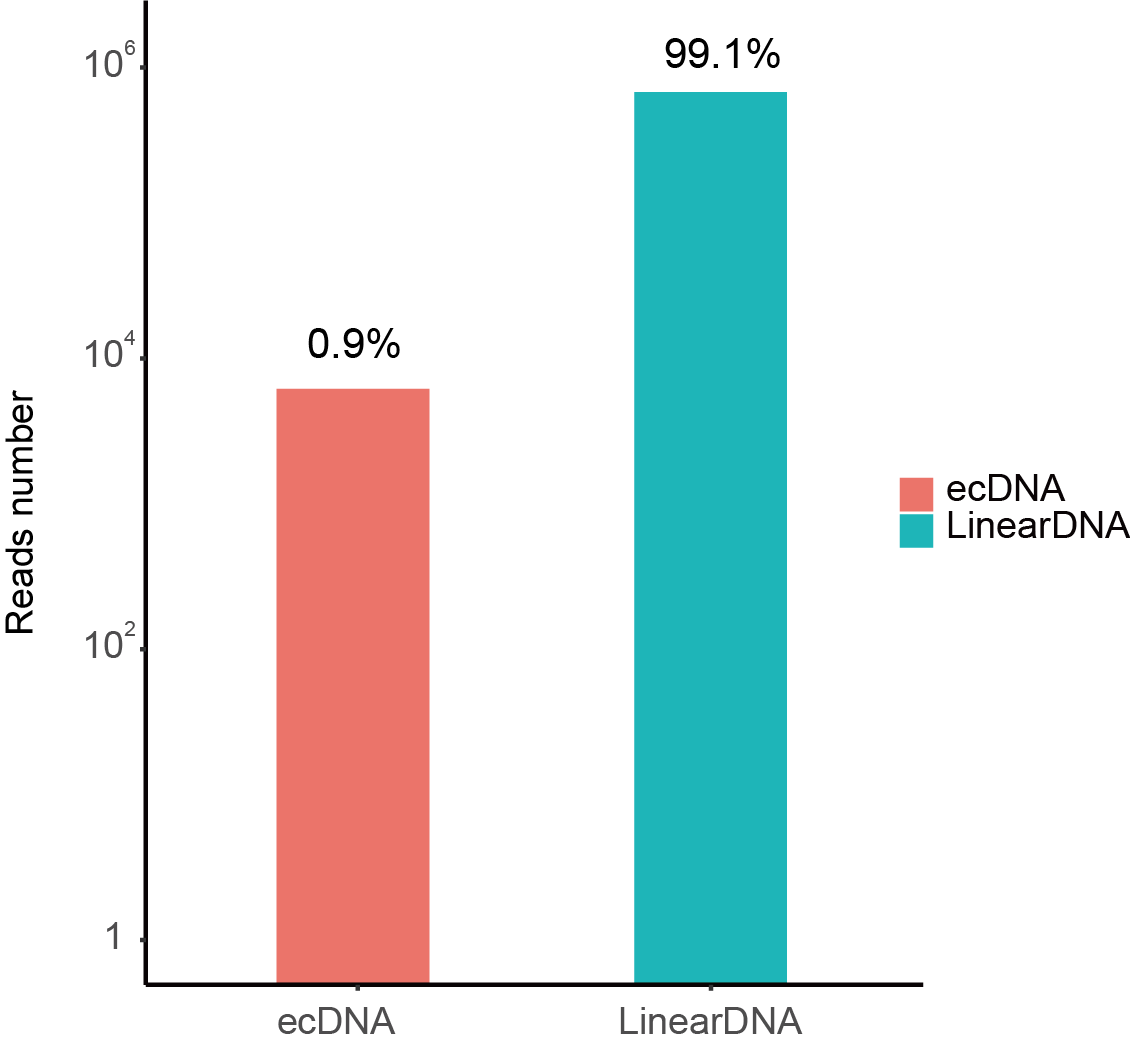

**Supplemental Figure 3. The counts of reads identified as ecDNAs or linear DNAs.**

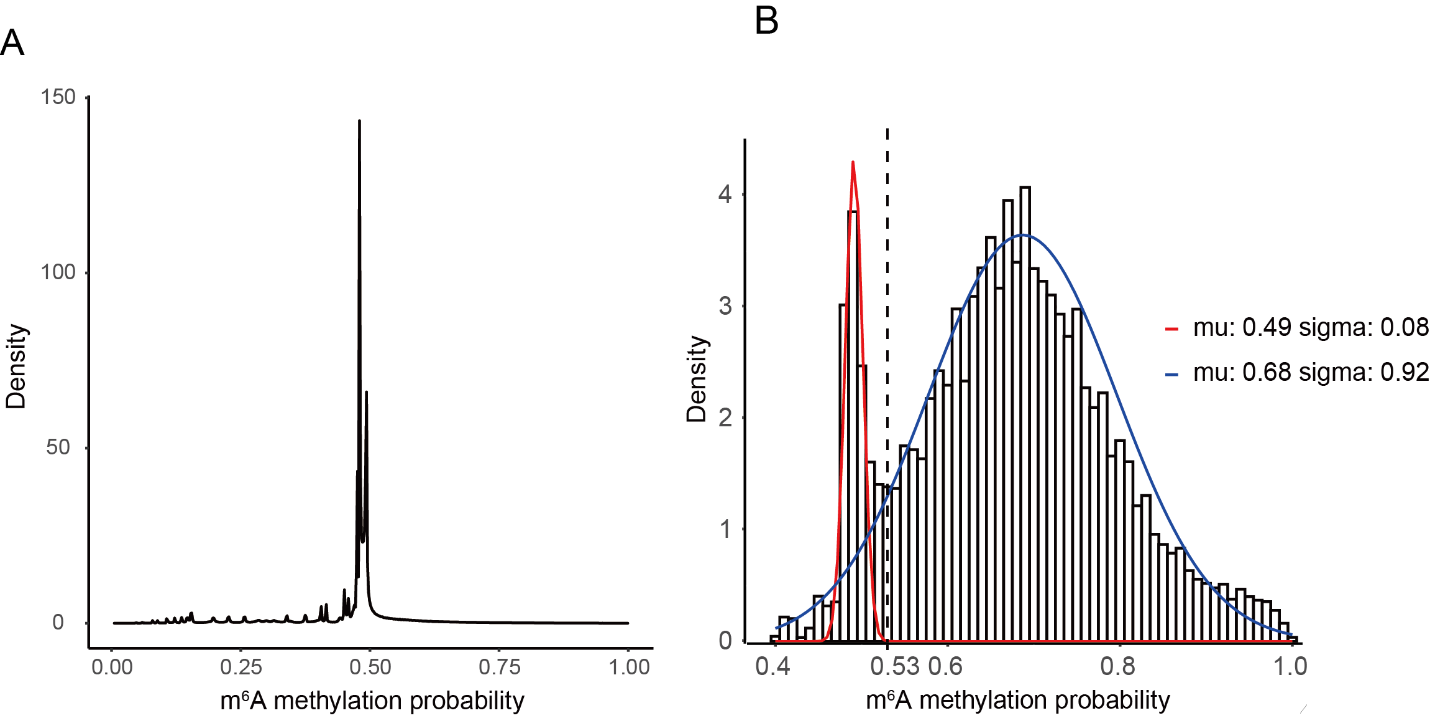

**Supplemental Figure 4. The m6A possibility distribution of the treated and non-treated samples.** The signal data were transformed to the m6A sites with possibility on each site. The m6A possibility distribution of the non-treated negative sample (A), which has no detectable m6A, was below 0.52. The two peaks (0.5,0.7) could be classified as positive sites and negative sites in EcoGII treated samples (B). The cut-off value was set as 0.53, and the m6A calling specificity and sensitivity is 0.99 and 0.92.

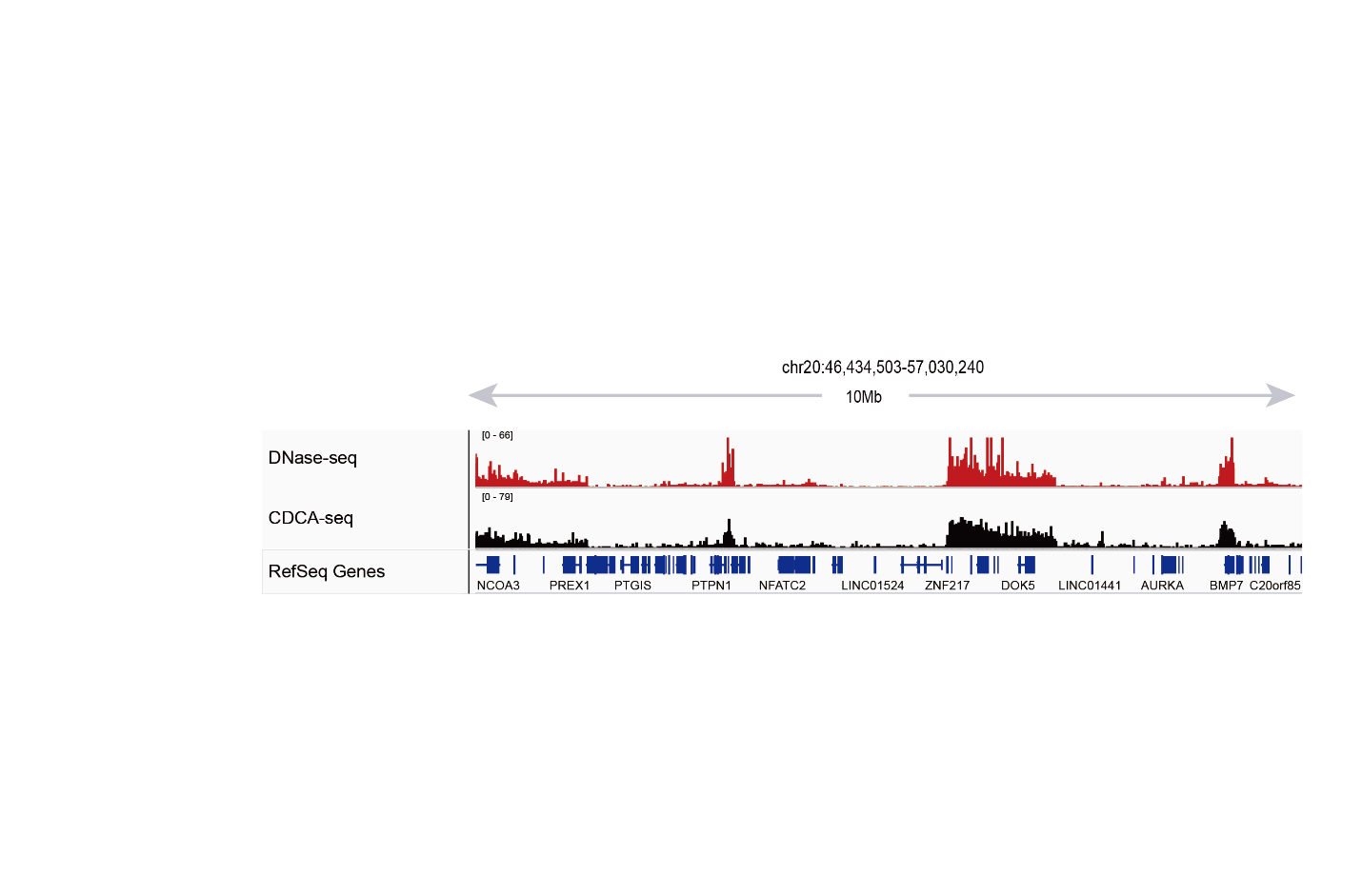

**Supplemental Figure 5. Large aggregate CCDA-seq signal enrichments match closely with DNase-seq accessibility peaks.** (Chr20:46,434,503-57,030,240)

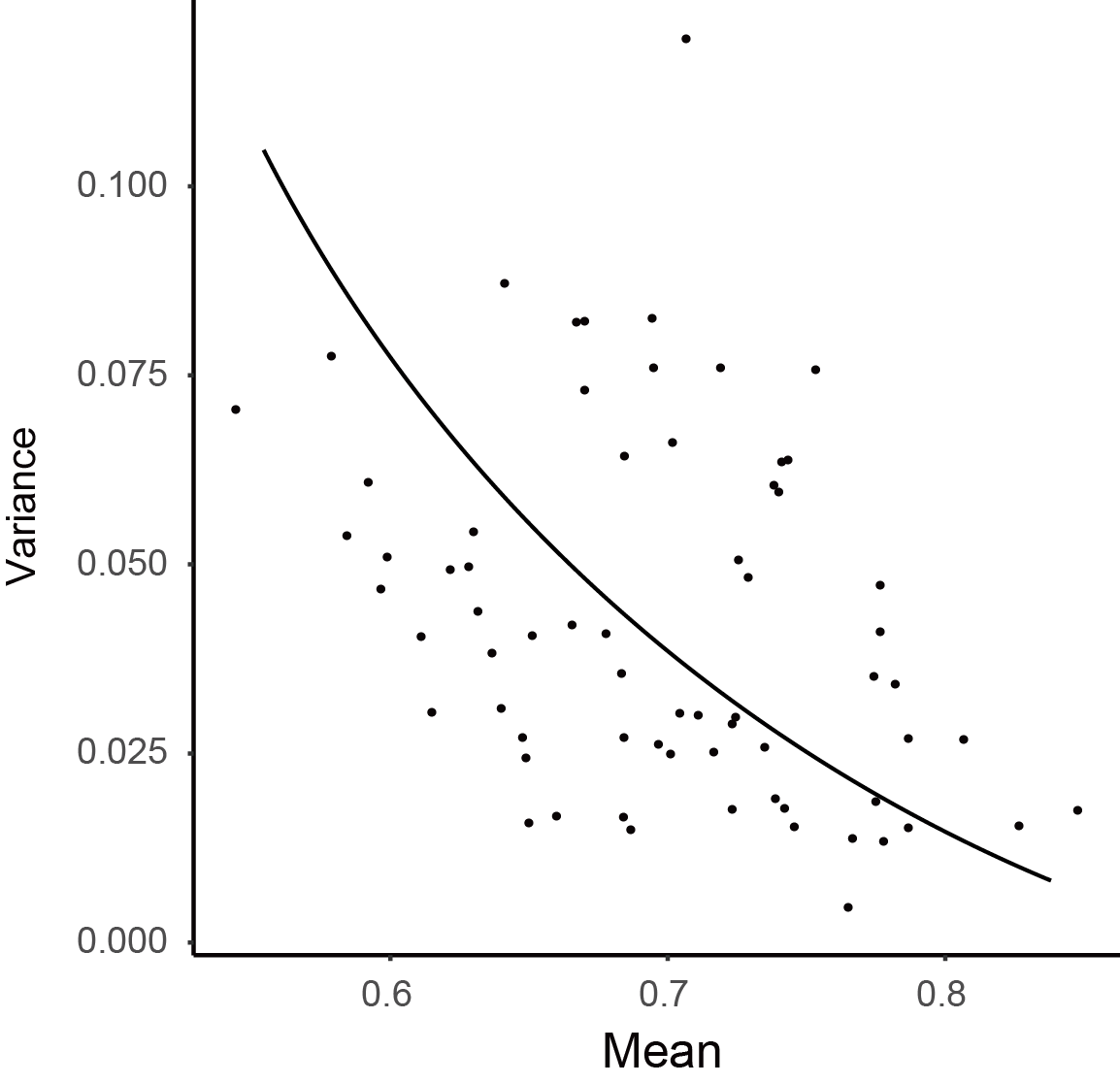

**Supplemental Figure 6. The m6A methylation deviation is related to the average m6A methylation.** The genome was sized into 50bp bins. The average m6A methylation was calculated as (total m6A in all covered reads)/(total adenosine in all covered reads). The deviation was represented by the bin methylation deviation, which aggregates the methylation in each bin.

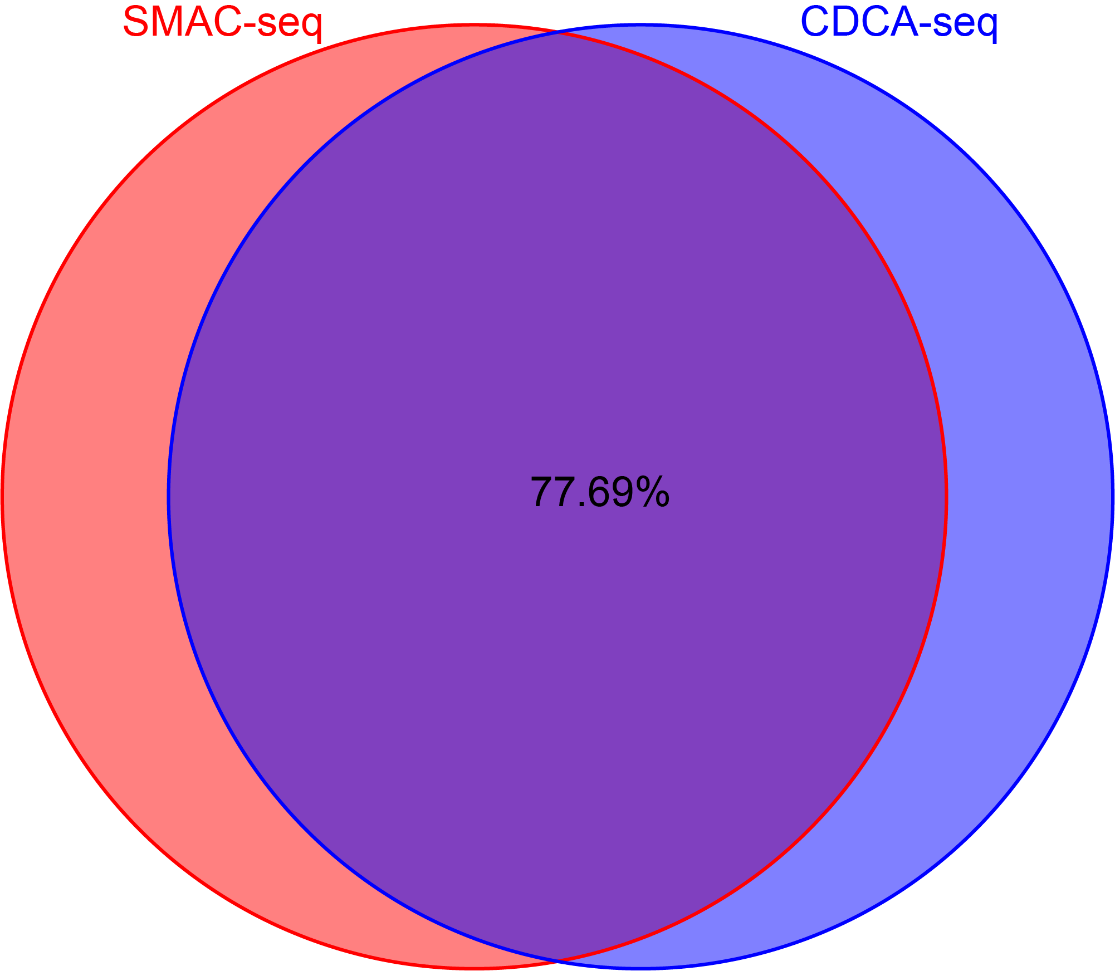

**Supplemental Figure 7. The correlation between two sample replicates with and without exonuclease treatment.** The methylated bins (methylated count>2, bin=50bp) are 77.69% overlapped between non-exonuclease-digested SMAC-seq and exonuclease-digested CCDA-seq.

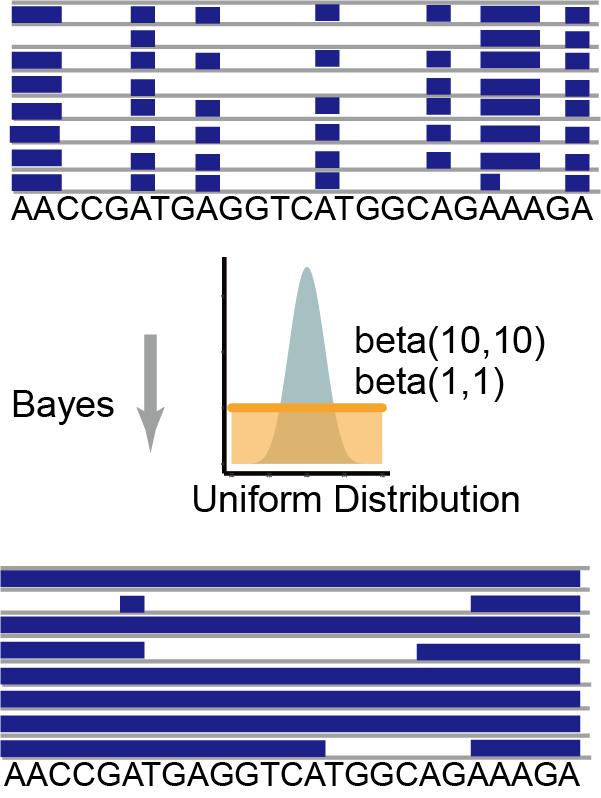

**Supplemental Figure 8. Outlines of the algorithm to configure the methylation level in genome resolution and single-molecule resolution.** In genome resolution, the methylation fraction represents the average methylation fraction of multiple reads covered in the regions. In the single molecular resolution, we adopted a Bayesian procedure to aggregate methylation probabilities and derived the accurate single-molecule accessibility calls over 50bp windows.

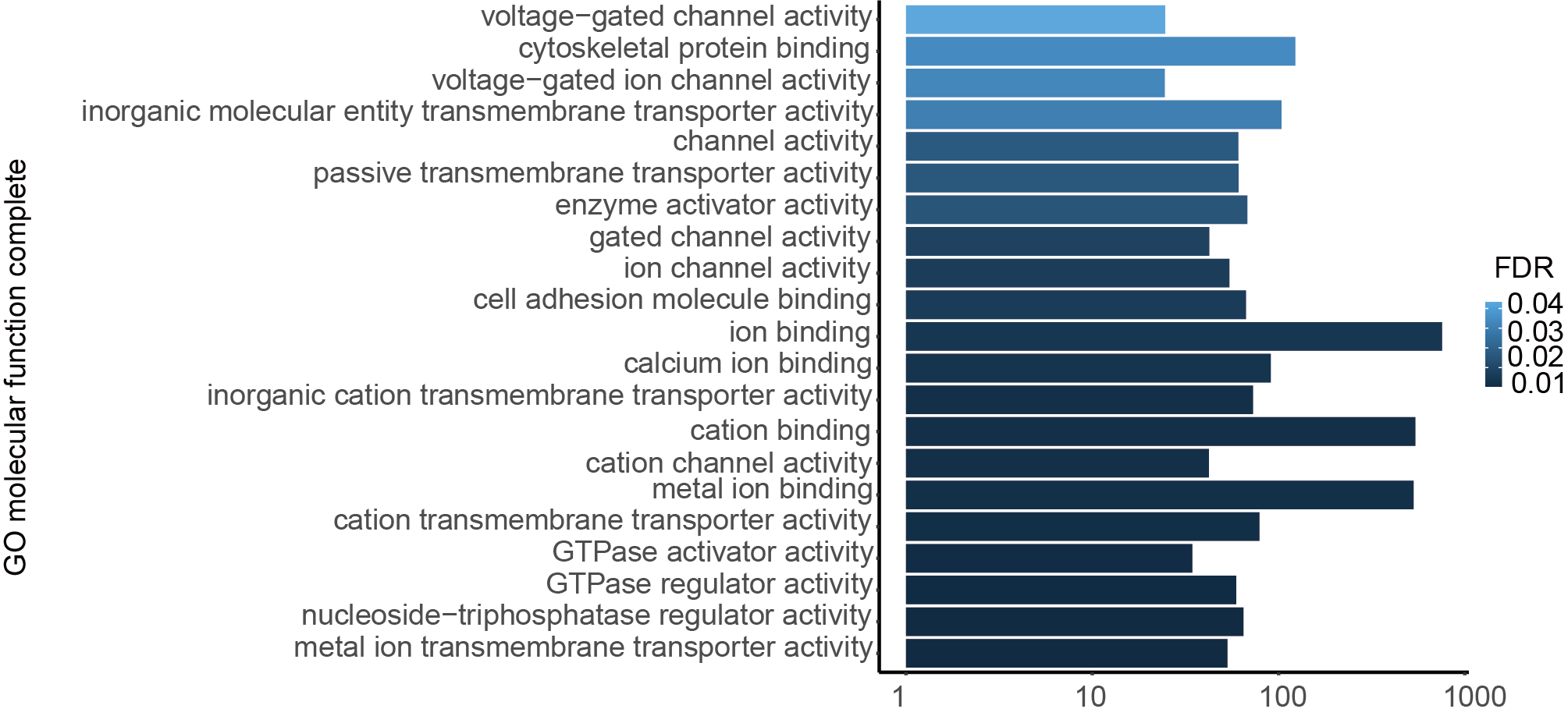

**Supplemental Figure 9. Gene ontology analysis of these ecDNA carried genes.** (<http://geneontology.org/>)

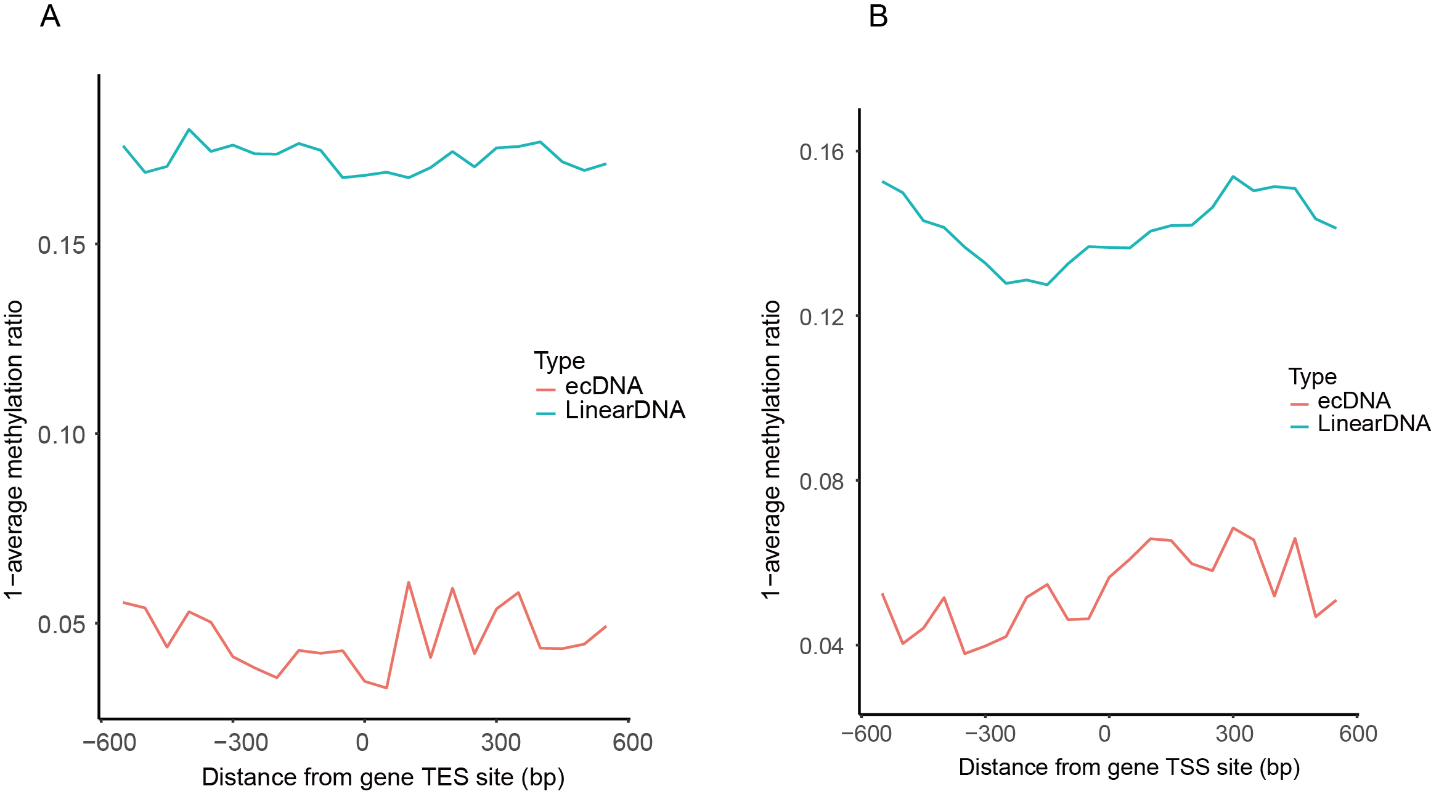

**Supplemental Figure 10. Average CCDA-seq profile around TSSs and TES of group II gene.** The group II genes have more open chromatin structure on ecDNA than linear DNA (Figure 2C).

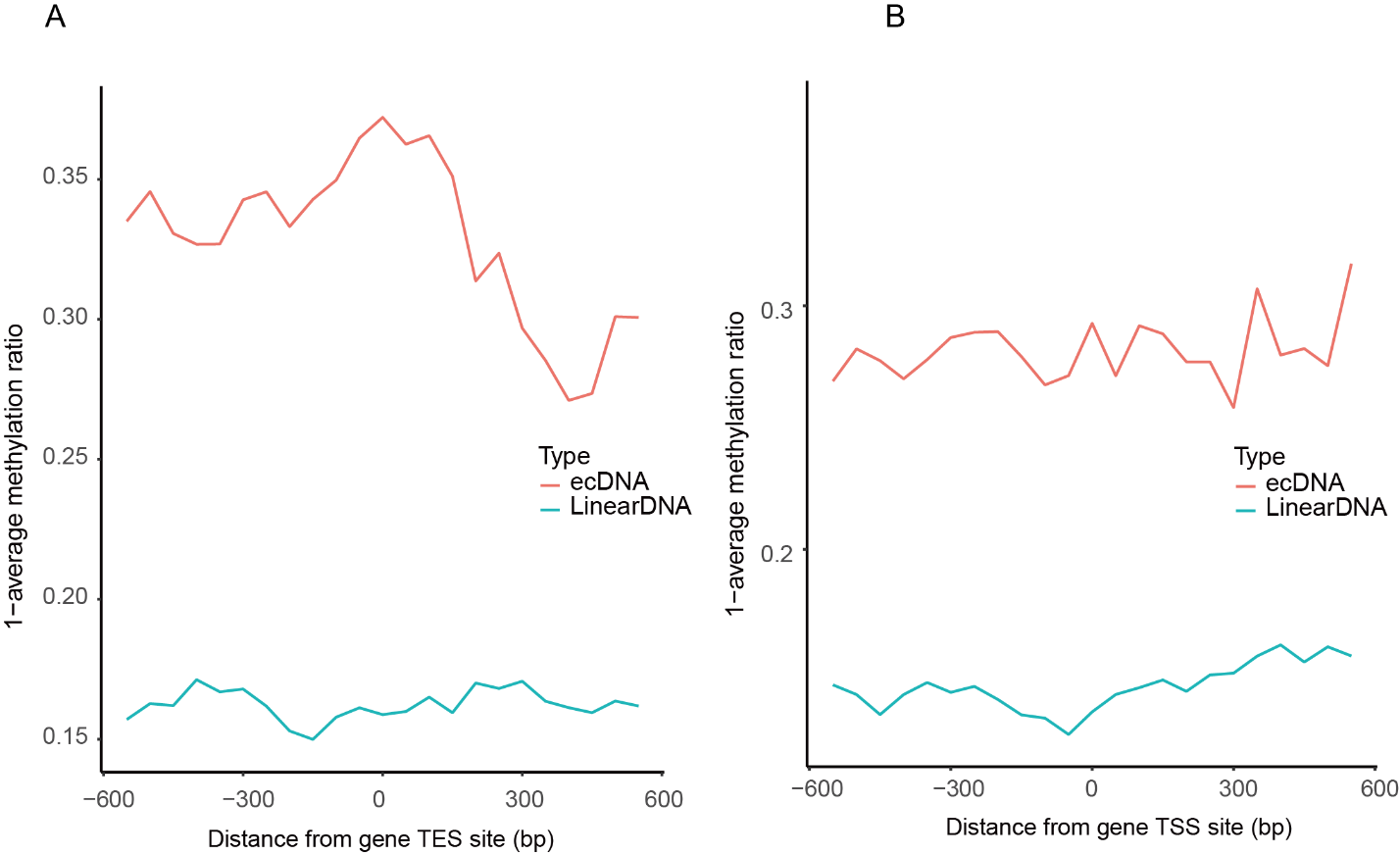

**Supplemental Figure 11. Average CCDA-seq profile around TSSs and TES of group I gene.** The group I genes have more open chromatin structure on linear DNA than ecDNA (Figure 2C).

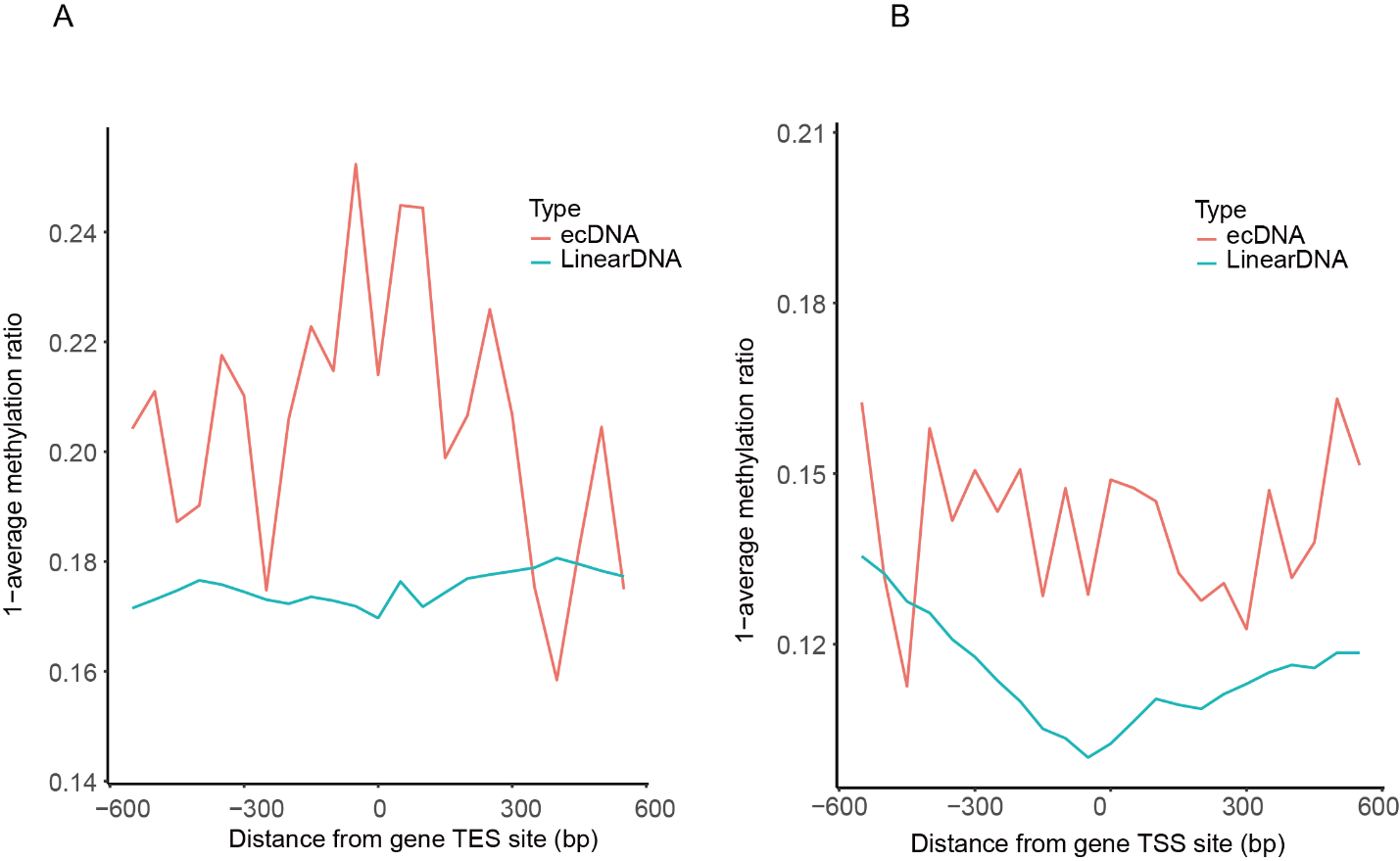

**Supplemental Figure 12. Average CCDA-seq profile around TSSs of genes with high expression level.** The top quantile defines the high expression genes in RNA-seq.

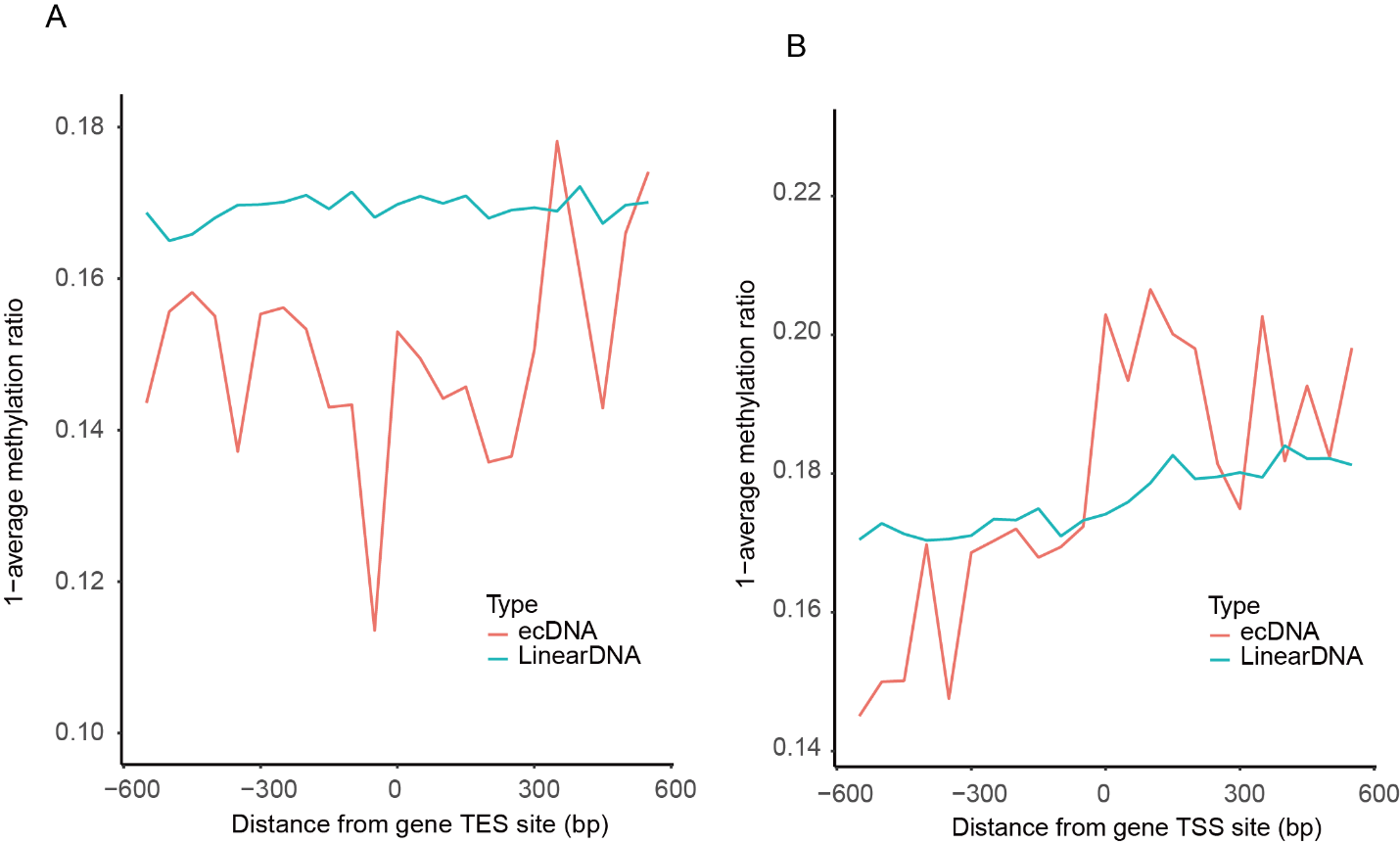

**Supplemental Figure 13. Average CCDA-seq profile around TSSs of genes with low expression level.** The bottom quantile defines the low expression genes in RNA-seq.

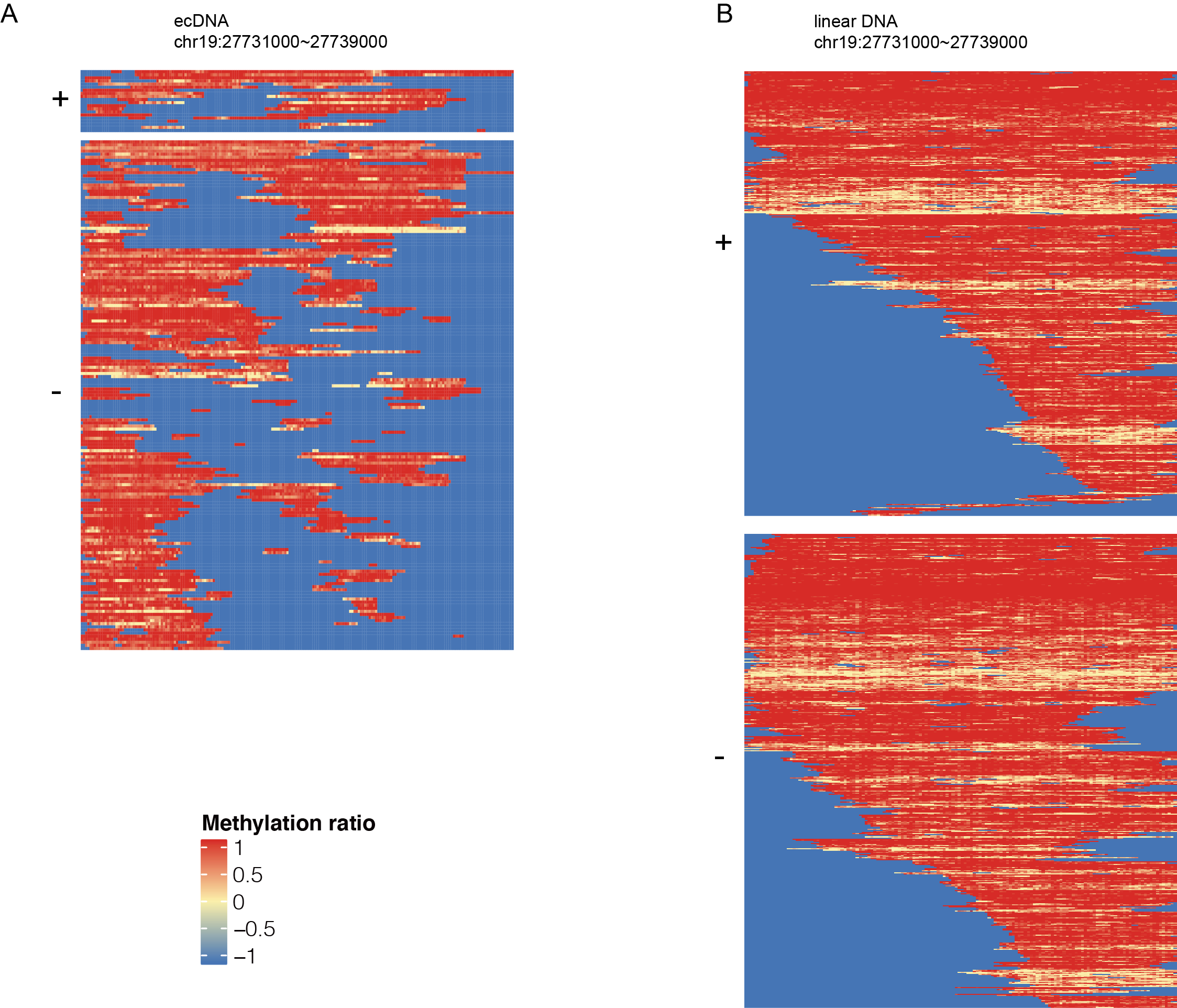

**Supplemental Figure 14. CCDA-seq reveal the distribution of alternative chromatin state in single molecular resolution.** A. Shown are all reads (+/-) covering ecDNA regions. B. Shown are all reads (+/-) covering the linear DNA areas.

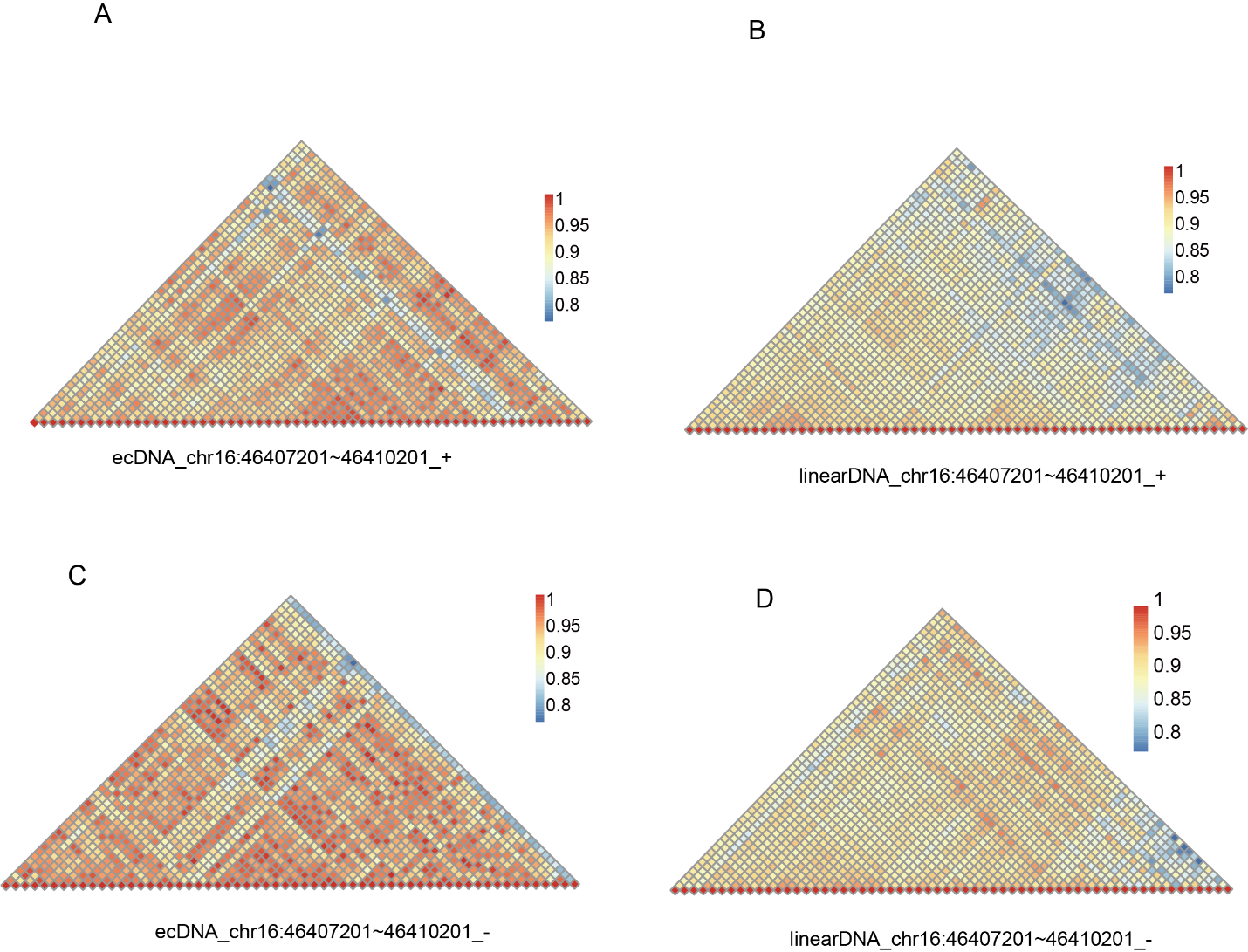

**Supplemental Figure 15. Chromatin co-accessibility profiles for the chr16:46407201~46410201 show correlation and anticorrelation in ecDNA and linear DNA.** (A) Chromatin co-accessibility profiles for the chr16:46407201~46410201 show correlation and anticorrelation on ecDNA positive strand. (B) Chromatin co-accessibility profiles for the chr16:46407201~46410201 show correlation and anticorrelation on linear DNA positive strand. (C) Chromatin co-accessibility profiles for the chr16:46407201~46410201 show correlation and anticorrelation on ecDNA negative strand. (D) Chromatin co-accessibility profiles for the chr16:46407201~46410201 show correlation and anticorrelation on linear DNA negative strand.

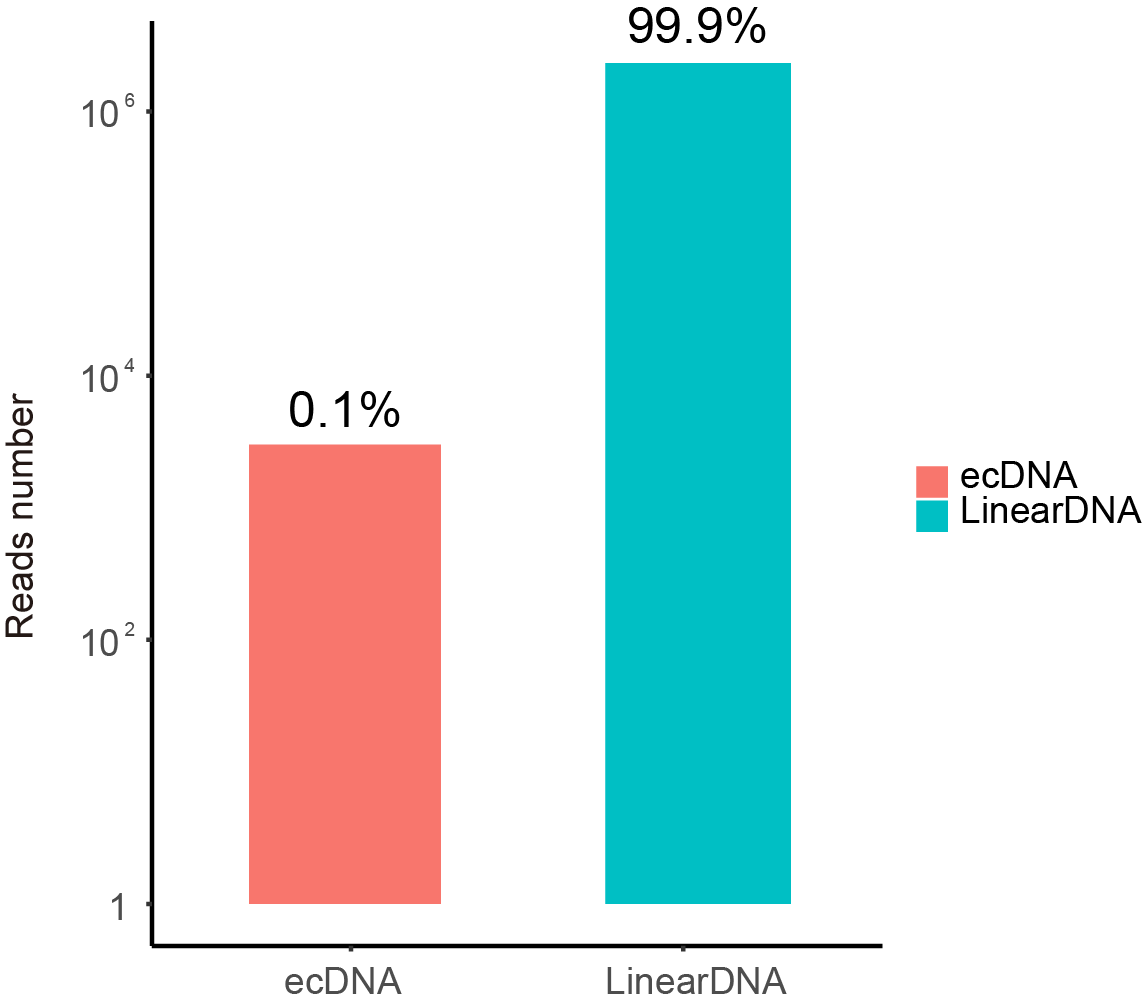

**Supplemental Figure 16. The counts of reads identified as ecDNAs or linear DNAs.** The sample is directly subjected to nanopore DNA sequencing without the exonuclease digestion to remove the linear DNA.

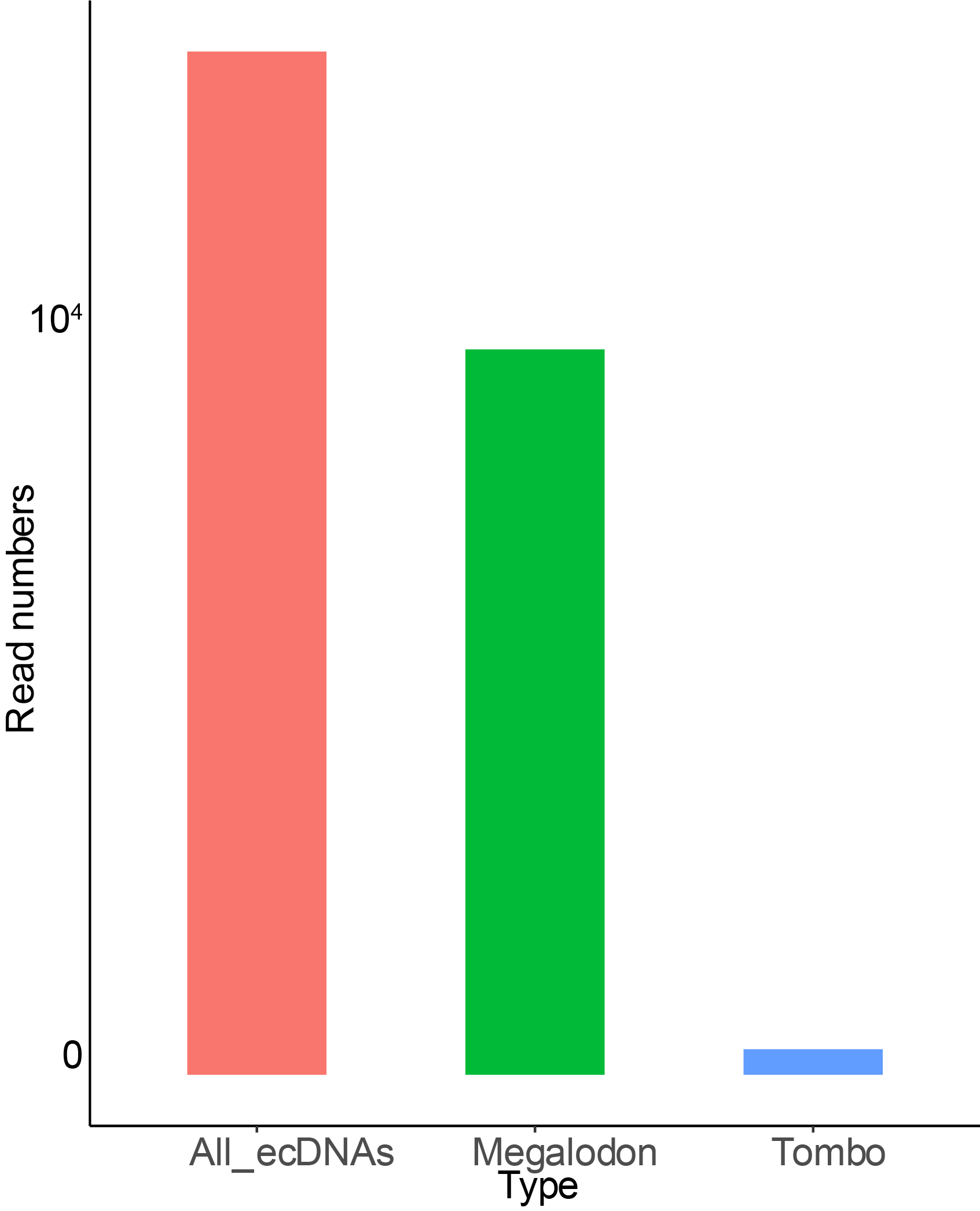

**Supplemental Figure 17. Comparing the megalodon and Tombo in ecDNA methylation calling.** All_ecDNAs indicate the ecDNAs identified by minimap2. All the detected ecDNAs are processed to call methylation by Megalodon or Tombo.

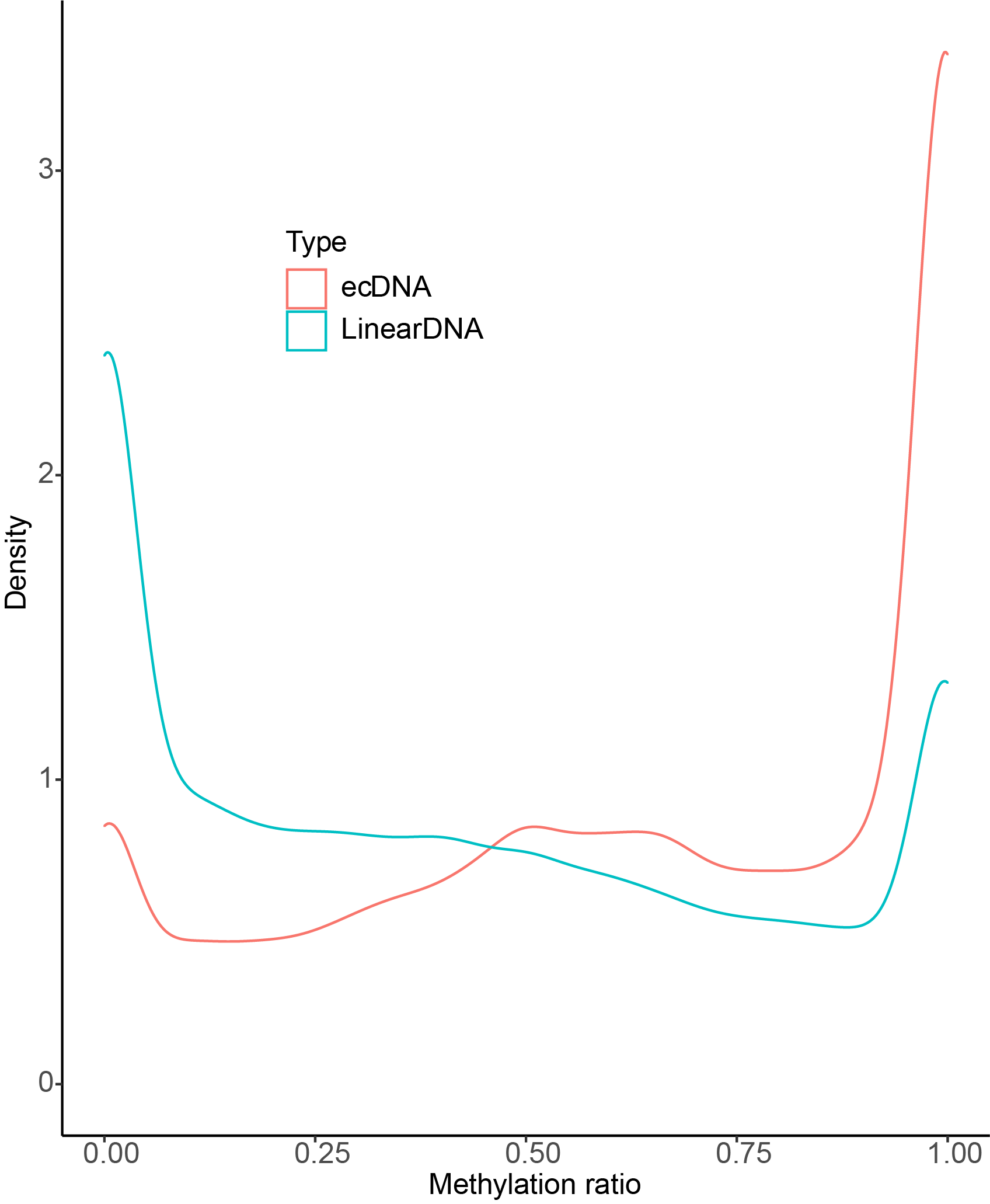

**Supplemental Figure 18. The density distribution of the methylation ratio on ecDNAs and linear DNAs in non-exonuclease digested sample**

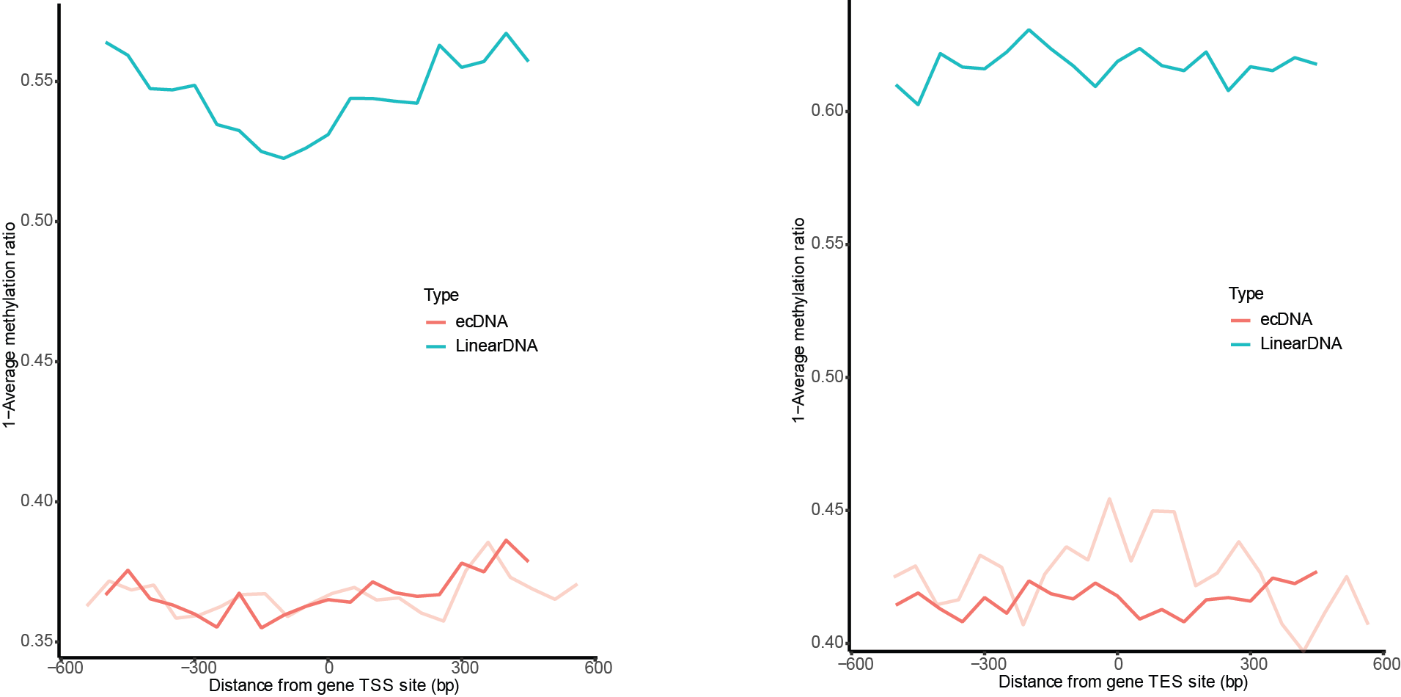

**Supplemental Figure 19. Average CCDA-seq profile (no linear DNA digestion) around transcription start site and transcription end site on ecDNAs and linear DNAs.** The sample, not digested by exonuclease, showed the lower average methylation on both eccDNAs and linear DNAs. The light red indicated the nucleosome occupancy in the CCDA-seq with exonuclease digestion. Comparison showed the approximately similar trend between the non-digested sample and digested sample. The difference may be caused by the distinct ecDNA coverage in two samples. The exonuclease treatment should not bias our analysis. (aggregated over 50-bp windows sliding every 5 bp; the sequencing depth is normalized for ecDNA and linear DNA;see Methods for details)

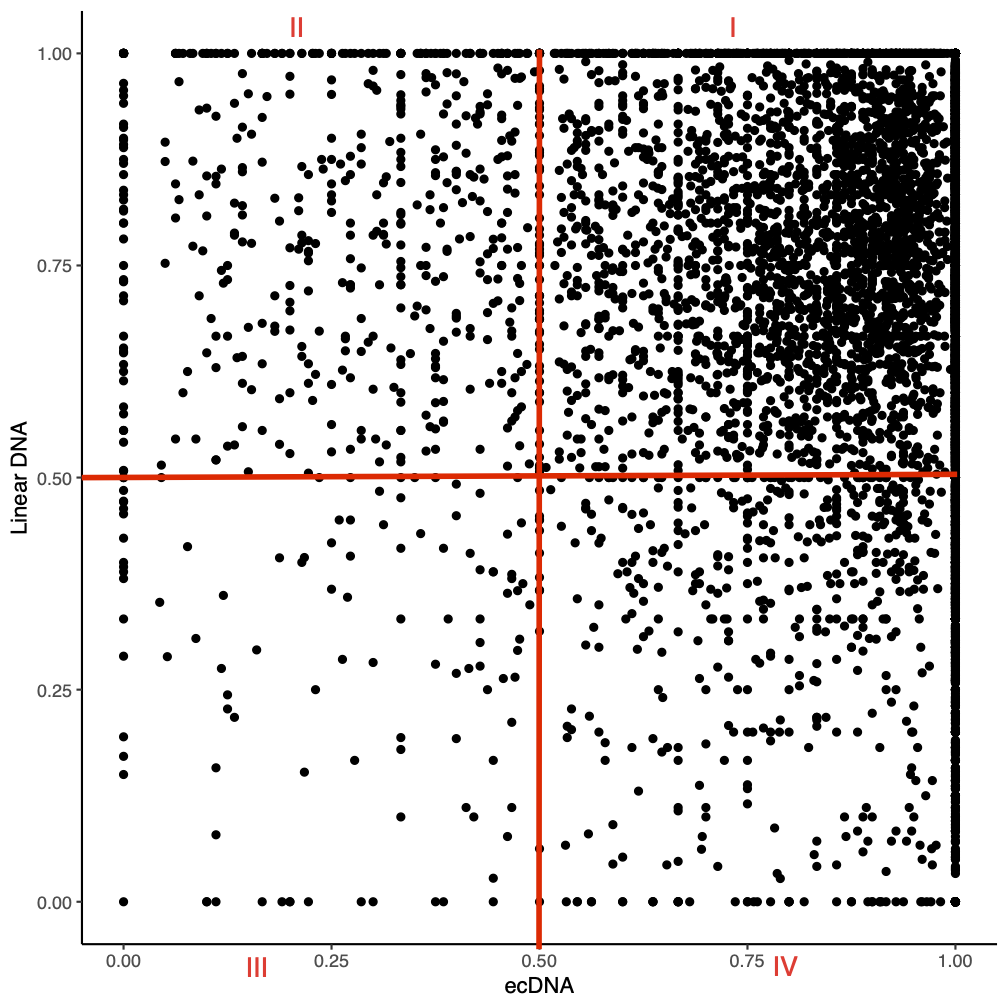

**Supplemental Figure 20. Pairwise scatter plot of the average bin methylation between ecDNAs and linear DNAs.** The genome is sized into 50bp bins. The methylation in each bin is the methylation ratio average of covering reads. The bins were classified into four groups: Type I- the regions are highly accessible in both linear DNA and ecDNAs; Type II- the linear DNA regions are less accessible than the ecDNA regions; Type III- the regions are not accessible in both linear DNA and ecDNAs; Type IV- the ecDNA regions have more open chromatin than the linear areas.

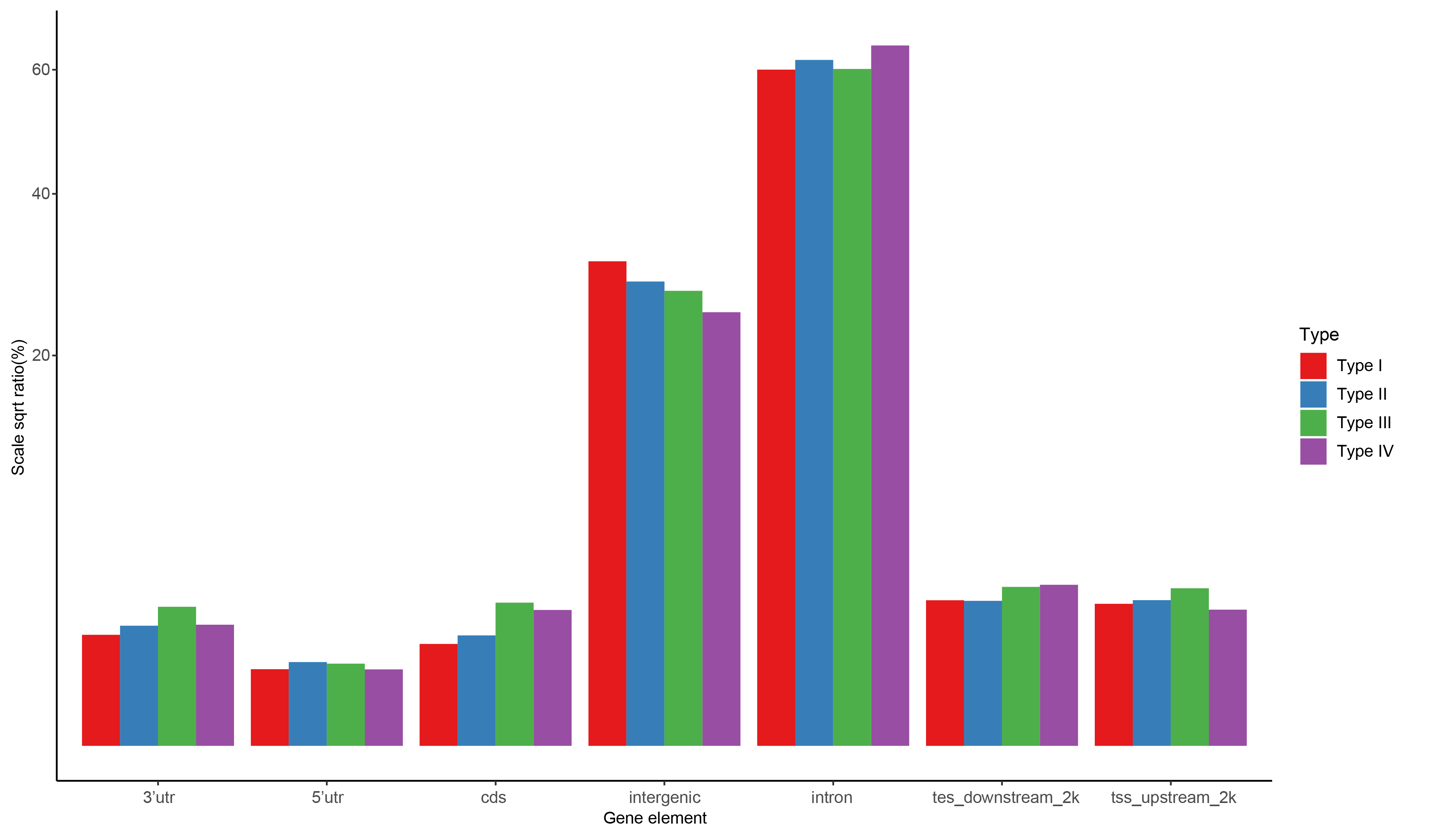

**Supplemental Figure 21. The bins distribution among the gene elements.** The four types of regions are defined in supplemental figure 18.
